# High-resolution synchrotron imaging studies of intact fresh roots reveal soil bacteria promoted bioremediation and bio-fortification

**DOI:** 10.1101/767558

**Authors:** Hanna Help, Merja Lusa, Ari-Pekka Honkanen, Ana Diaz, Mirko Holler, Murielle Salomé, Peter Cloetens, Henrik Mäkinen, Simo Huotari, Heikki Suhonen

## Abstract

Plant-microbe interactions can be utilized in bio-based processes such as bioremediation and biofortification, either to remove hazardous radionuclides and heavy metals from the soil, or to increase the accumulation of desired elements into crops to improve their quality. Optimizing such elegant plant-microbe interactions requires detailed understanding of the chemical element compositions of fresh plant tissues at cellular organelle resolution. However, such analyses remain challenging because conventional methods lack the required spatial resolution, contrast or sensitivity. Using a novel combination of nanoscale-resolution 3D cryogenic synchrotron-light ptychography, holotomography and fluorescence tomography, we show how soil bacteria interact with *Arabidopsis thaliana* and promote the uptake of various metals. Co-cultivation with *Pseudomonas* sp. strain T5-6-I alters root anatomy and increases levels of selenium (Se), iron (Fe) and other micronutrients in roots. Our approach highlights the interaction of plants and microbes in bioremediation and biofortification on the subcellular level.

---

Soilborne bacteria can modify soil chemical landscapes altering the mobility and uptake rates of various chemical elements, including hazardous radionuclides and heavy metals, but also macro- and micro-nutrients into plant tissues (Ojuederie *et al.*, 2018, Desai *et al.*, 2019). For tailoring plant-microbe interactions to improve the efficiency of bio-based processes like bioremediation and biofortification, conclusive data on chemical element composition from intact fresh plant tissues at cellular organelle resolution is required. Currently, however, there is a lack of quantitative knowledge on the ultimate fate of accumulated elements in plants on a subcellular level. This is unfortunate, as understanding in which tissue and cellular organelle various chemical elements and species accumulate would help increase their uptake and translocation rates e.g. by modifying tissue and organelle specific transporter activity via genetic engineering.

Selenium (Se) is an essential micronutrient for mammals, but in high-valence states it is toxic at elevated concentrations. In alkaline agricultural soils, selenium mostly exists as selenate (Se(VI)O_4_^2−^) whereas in acidic forest soils it exists as selenite (Se(IV)O_3_^2−^). The environmental releases of Se in these forms is potentially hazardous (Gupta & Gupta, 2016). Selenate and selenite differ in their absorption and mobility within the plant (Li *et al.*, 2008). As the environmental mobility and biological effects of Se are mainly dictated by its chemical speciation, it is interesting that the chemical speciation can be affected by some soilborne bacteria, including *Pseudomonas* sp. strain T5-6-I, via reduction and oxidation (redox) reactions. T5-6-I forms intracellular aggregates of reduced Se^0^ when incubated in Se(IV)O_3_^2−^ containing solutions, and co-cultivation with plants enhances Se(IV) uptake and accumulation *in planta* (Lusa *et al.*, 2015, 2017 and 2019). Indirect evidence suggests that in plants Se accumulates in cellular vacuoles (Gupta & Gupta, 2016, Santa Cruz *et al.*, 2017) as it does in human tissues (Bodnar *et al.*, 2017). Transmission electron microscopy (TEM) (Durán *et al.*, 2018, Lusa *et al.*, 2019) has revealed aggregate formation in plant cell compartments after Se treatments. However, it was not concluded if the observed aggregates were selenium. While direct chemical element localization has been done from fresh plant roots (Wang *et al.*, 2013, Hu *et al.*, 2014, Zhao *et al.*, 2014, Krajcarová *et al.*, 2017) and seeds (Rodrigues *et al.*, 2018) using synchrotron radiation x-ray fluorescence microscopy, the resolution has not been sufficient to determine subcellular element concentrations. Using sectioning techniques and 2D imaging methods, such as nano-secondary ion mass spectrometry (nano SIMS), laser ablation-inductively coupled plasma-mass spectrometry (LA-ICP-MS), or ColorEM (electron microscope) one could potentially reach desired resolution and sensitivity (Ondrasek *et al.*, 2019, Pirozzi *et al.*, 2018). However, these techniques can only show small fraction of the total volume and are subject to artefacts from the sectioning process. Therefore, the question whether plants form intracellular vacuole localized Se granules in their roots, similarly to *Pseudomonas* sp. strain or humans, or if Se is deposited elsewhere in plant tissues (e.g. cell walls) has not been answered conclusively.

We established the role of bacteria on the accumulation and distribution of Se in the elongation zone of roots, the key uptake sites for water and nutrients, by using multimodal synchrotron nanotomography on fresh Arabidopsis thaliana root material. We did this by imaging frozen native state specimens cultivated in Se(IV) and Se(IV) + *Pseudomonas* sp. strain T5-6-I supplemented growth media with three state-of-the-art synchrotron x-ray imaging techniques: X-ray ptychography (PXCT) (Dierolf *et al.*, 2010) and nanoholotomography (XNH) (da Silva et al. 2018), both giving high resolution 3D images of the electron density, as well as x-ray nanofluorescence microscopy (XRNF) giving quantitative, high resolution 2D and 3D images of trace element distributions in the plants. These techniques gave subcellular organelle resolution, allowing to address whether the aggregates reported with EM studies (Durán *et al.*, 2018, Lusa *et al.*, 2019) are indeed composed of Se.

Our findings show that, even though Se levels varied between different treatments, tissue profiles of Se and Se+bacteria treated samples were similar. Most importantly; soil bacteria appeared to promote Se accumulation in young *Arabidopsis thaliana* lateral root tips. Using tissue segmentation analysis and subcellular measurements, we were able to determine that Se concentrates predominantly to cytoplasmic domains, cell walls and vacuolic membranes in roots. Based on our results we were able to conclude that Se does form aggregates in *Arabidopsis thaliana* plants, however not in a similar manner as has been reported with bacterial or mammalian cells, where Se aggregates predominantly in vacuolic compartments. Interestingly, bacterial co-culturing also promoted accumulation of Fe and other micronutrients into root tissues. Fe was predominantly localized in cell walls showing different localization pattern than Se. The amount of Fe in bacteria co-cultured root tissues exceeded Se levels by a magnitude and could partially explain the observed increase in root cell wall density. From an agricultural point, soil bacteria promoted Fe biofortification would be a cost-effective way to improve the nutritional value of crops growing in micronutrient poor soils.

## RESULTS

### Root tissue densities and physical features differ between treatments

Overall, in all imaged lateral root samples (Figure 1 A), cell walls had the highest electron density, followed by vacuole membranes, cytoplasm (including endoplasmic reticulum and different cell organelles like mitochondria) and vacuolic fluid (least dense) (Figure 2, Extended Figure 1). Interestingly, we found that bacteria increased cell wall densities by >7% compared to control samples, while the cytoplasm and vacuolic fluid densities were similar in all samples irrespective of treatment (Table 1). The vacuolic membrane densities of the Se+bacteria treated root were ~16% higher than in Se-alone supplemented sample. This difference is large enough to be directly seen in electron density visualizations (Figure 2, panels I and L).

**Figure 1:**
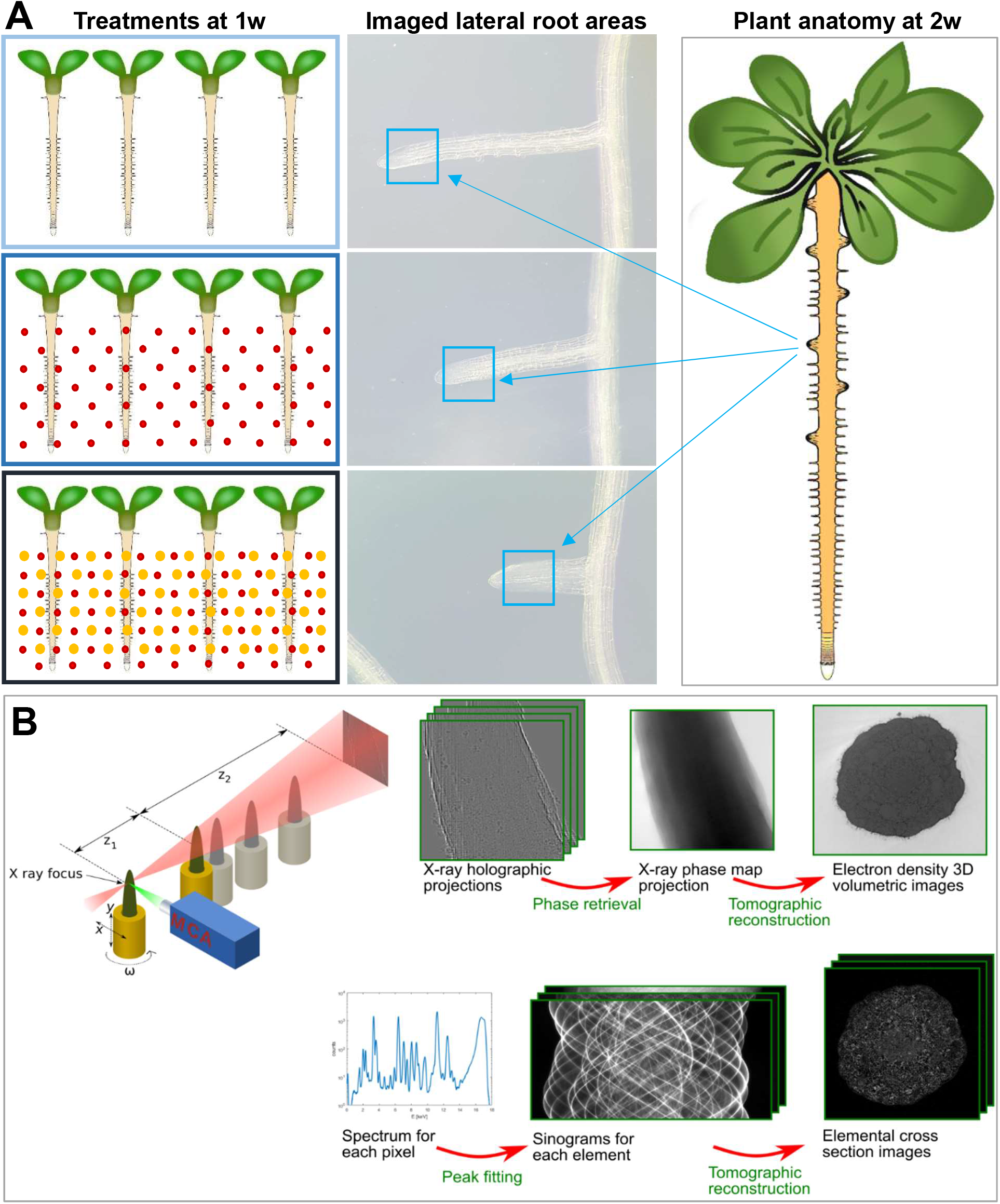
Plant material and visualization of the ID16A imaging protocol. A) Arabidopsis seedlings were grown in 3 different conditions: Normal 1% suc 1xMS media (left, top), 1 w on normal media + 1 w on Se supplemented media (left, middle) and 1 w on normal media + 1 w on Se supplemented media with bacterial co-culturing (left, bottom). Roots of same age were collected few days after emergence (middle panels, note the growing conditions affected root elongation rates) from ~2w old plants in vegetative growing phase showing few rosette leaves (illustration on the right). Collected roots were mounted in cryogenic imaging holders. The imaged areas are shown in small white boxes in middle panes (middle-top, control root; middle-middle Se supplemented; middle-bottom, Se+bacteria cultured root). B) illustration of the ID16A imaging protocol: 2D XNH holotomographic projection images taken at different distance from X-ray focus were used to retrieve phase map projection images. These 2D phase images were then reconstructed to 3D volumes. XRNF measurements were conducted by placing the sample in the X-ray focus, and then raster scanning the sample pixel by pixel to produce 2D maps and sinograms. The XRNF 2D slices and 3D stacks (calculated from sinograms) were located at the center of the XNH field of view, at 115 μm distance from the root tips.

**Figure 2:**
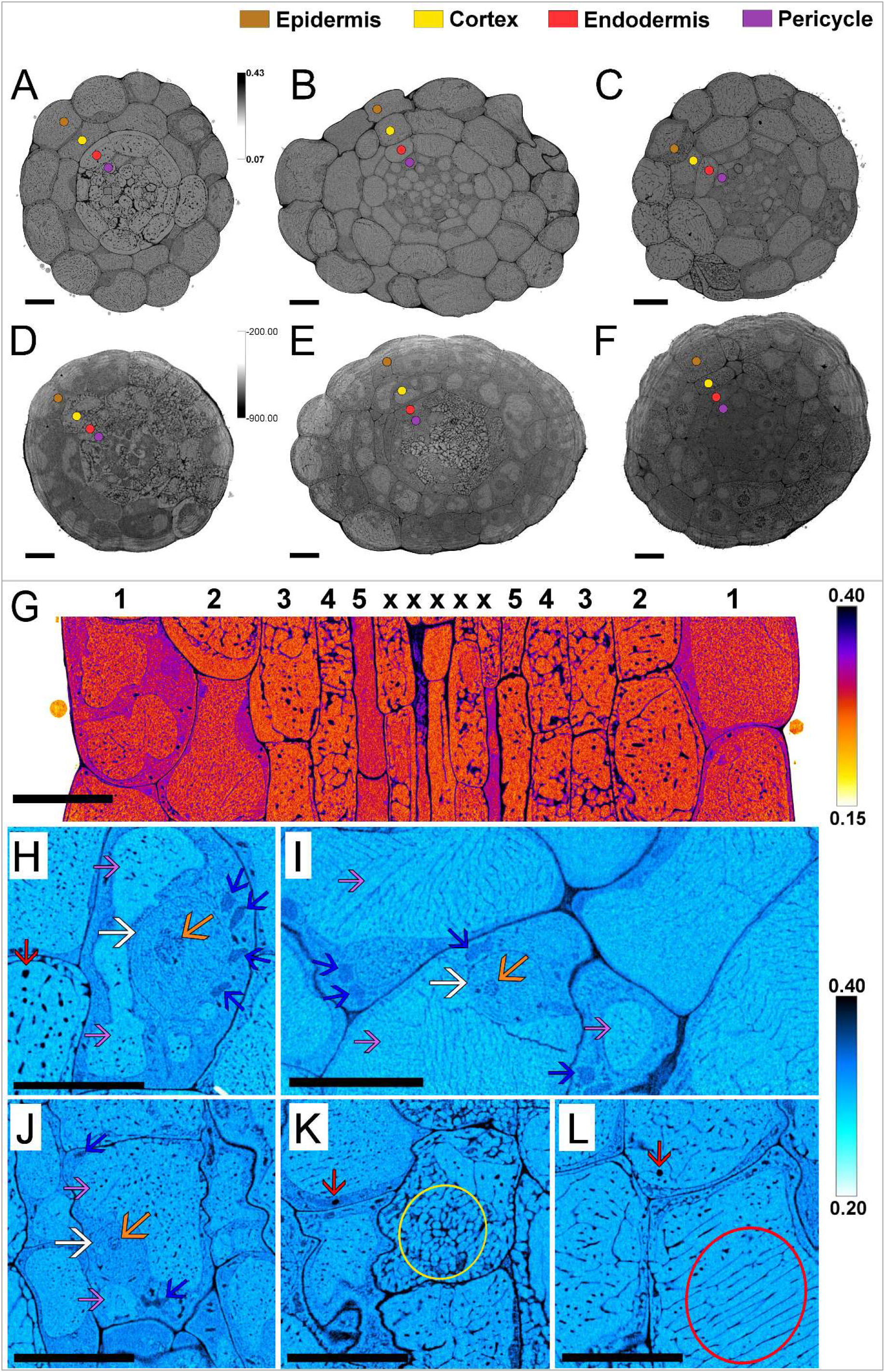
Anatomical comparisons and details of PXCT and XNH imaged roots. Electron density cross section images from PXCT (A – C) and XNH (D - F) reconstructions. A) control root, B) Se supplemented root, and C) Se+bacteria supplemented root. D) control root, E) Se supplemented root, and F) Se+bacteria supplemented root. Overall the imaged roots appear to be anatomically very similar within each treatment (A and D, control roots) (B and E, Se supplemented roots) (C and F, Se + bacteria supplemented roots). Identities of different cell layers are indicated with colored dots in the greyscale electron density images; epidermis (brown), cortex (yellow), endodermis (red) and pericycle (purple). An electron density signal intensity-based color image illustrates different cell types in a longitudinal cross section of control root: 1) epidermis, 2) cortex, 3) endodermis, 4) pericycle, 5) phloem sieve element (cleared from cytoplasmic organelles), x) procambium/ metaphloem cells. H - L) Close up images of heat-map colored control root (H), Se supplemented root (I) and Se+bacteria supplemented root (J, K and L). These close ups of epidermal and cortical cells nicely illustrate that cryogenic 3D PXCT imaging allows inspection of density features in very fine detail, even at the level of subcellular organelles. Marked in the image are various subcellular and nuclear compartments: nuclear envelopes (white arrows) and nucleoli (orange arrows). Nuclear cavities and chromatin are within the nuclear envelope. The nucleus is surrounded by mitochondria (dark blue arrows), which appear denser that the surrounding cytoplasmic fuid and vacuolic “fluid” within the expanding vacuolic compartments (lilac arrows). Red arrows point to dense aggregates. Yellow circle in panel K illustrates high density cytoplasmic membranes (ER), and red circle illustrates dense vacuolic membranes, All scale bars 10μm.

**Table 1:** Overview of the cryo-PXCT measurement parameters. Overview of the cryo-PXCT measurement parameters. (*) The FSC estimation for id134 is unrealistically low due to the insufficient amount of projections in each half-tomogram, given the large diameter of this sample.

| Sample ID | e- density |  |  |  |
| --- | --- | --- | --- | --- |
|  | min | max | median | mean |
| control cell wall | 0.36 | 0.42 | **0.39** | **0.39** |
| control cytoplasm and organelles | 0.28 | 0.37 | 0.32 | 0.32 |
| control vacuole fluid | 0.26 | 0.34 | 0.30 | 0.30 |
| control vacuole membranes | 0.33 | 0.42 | 0.38 | 0.38 |
| Se cell wall | 0.31 | 0.43 | 0.38 | 0.38 |
| Se cytoplasm and organelles | 0.26 | 0.38 | 0.32 | 0.32 |
| Se vacuole fluid | 0.25 | 0.35 | 0.30 | 0.30 |
| Se vacuole membranes | 0.28 | 0.40 | **0.34** | **0.34** |
| Se+bact cell wall | 0.33 | 0.47 | **0.42** | **0.42** |
| Se+bact cytoplasm and organelles | 0.28 | 0.40 | 0.33 | 0.33 |
| Se+bact vacuole fluid | 0.27 | 0.34 | 0.31 | 0.31 |
| Se+bact vacuole membranes | 0.32 | 0.45 | **0.40** | **0.39** |

Presented values of subcellular organelle density minimum. maximum. median and mean from different treatments are averages. Each measured from n>30 areas around the sample. The size of the measured areas depended on the cell organelle size.

The ground tissues (the two outermost cell layers, epidermis and underlying cortex cells) of control roots (Figure 2 A and D) appeared denser than the underlying endodermal and stele cells. Se treated roots (Figure 2 B and E) did not exhibit obvious differences in tissue density profiles between the cell types. Se+bacteria supplemented roots exhibit higher density of endodermis, pericycle and the stele cells (vasculature) than in the control or Se supplemented plants. In addition, Se+bacteria supplemented roots exhibited pronounced cytoplasmic and vacuolic membrane features and numerous high-density aggregates, mainly in ground tissue layers (Figure 2, panels K and L). Apart from density features, 3D visualizations of the datasets revealed differences in cellular differentiation rates between control, Se supplemented and Se+bacteria supplemented plants (Figure 3). Compared to control root (Figure 3 A, D), Se treated roots exhibited enhanced vacuolization of cortical cells, but no other visible abnormalities (Figure 3 B, E). On the contrary, Se+bacteria supplemented roots exhibited both enhanced vacuolic expansion and abnormal cell division patterns (Figure 3, C and F). These aberrations in cellular division and differentiation rates appeared to correlate with the increased numbers of root hairs in *Arabidopsis* roots (Figure 3 panels G, H and I), similar to what was reported on kale (Lusa *et al.*, 2019).

**Figure 3:**
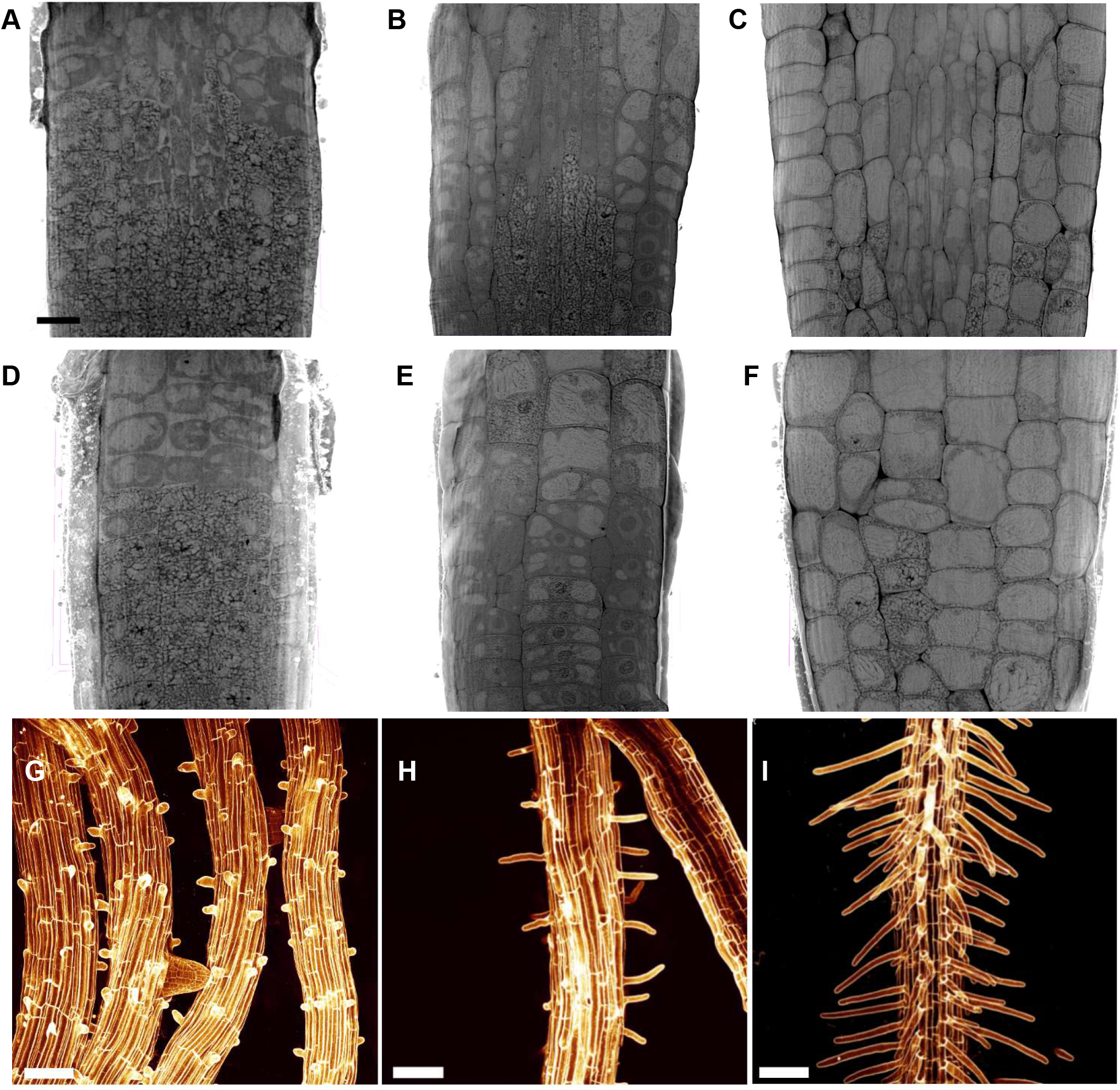
Meristem anatomical features and root hair densities. Meristem anatomy of control (A and D), Se (B and E) and Se + bacteria treated roots (C and F) differ from each other. Panels A, B and C represent longitudinal cross sections showing the vasculature, whereas panels D, E and F show ground tissue layers. Arabidopsis control root hairs in panel G, Se supplemented root in H, and Se+bacteria supplemented root in I were imaged using a confocal laser microscope. The root hair phenotypes are likely a combination resulting from selenium toxicity (with additive effects of other micronutrient accumulation,) and bacterial elicitor sensing and uptake (that mimic plant phytohormone, e.g. auxin signaling). Scale bar in panel A is 10μm and 50μm in panels G - I.

### Micronutrient accumulation differs between treatments

We found that not only the uptake of Se is modified by bacteria, but most notably that Fe uptake is promoted, and to smaller extent also that of Br and Cr (Figure 4, Extended Figure 2). The uptake or distribution of other detected elements as Al, Au, Ca, Cl, Cu, K, Mn, Ni, P, Pb and S was not modified by the presence of Se or bacteria. In Figure 4 the distinct cellular features of the XNH datasets are matched to XRNF-microscopy maps to visualize Se, Fe, Cr and Ca distribution (Figure 4). Top row of figure 4 illustrates the electron density slices that correspond to chemical element maps (1, control root; 2 and 3, Se supplemented roots; 4 and 5 Se+bacteria supplemented roots). As expected, control roots did not contain Se above the detection limit (Graph 1), while considerable levels of Se were detected in all root tissues of plants supplemented with Se. Overall, the tissue Se profiles of roots supplemented with Se-alone were comparable to Se+bacteria roots – however clear root-to-root variation in the overall levels of Se was seen between biological replicates of Se-alone and Se+bacteria treated samples (Graph 1). On tissue level, accumulation of Se appears quite even between intracellular and cell wall areas in Se and Se+bacteria supplemented roots (Graph 2). The Se+bacteria supplemented root #2 contained more Se than other measured samples. Interestingly, the Fe levels in control and Se supplemented roots were very low compared to Se+bacteria supplemented roots, which contained over 10-times more Fe than the other measured samples (Graphs 1 and 2). Unexpectedly, the levels of Fe of Se supplemented roots were even lower than in control root, with the fluorescent signal of Se supplemented roots localized mainly to the superficial cell facing growing medium (Figure 5, Graph 2).

**Figure 4:**
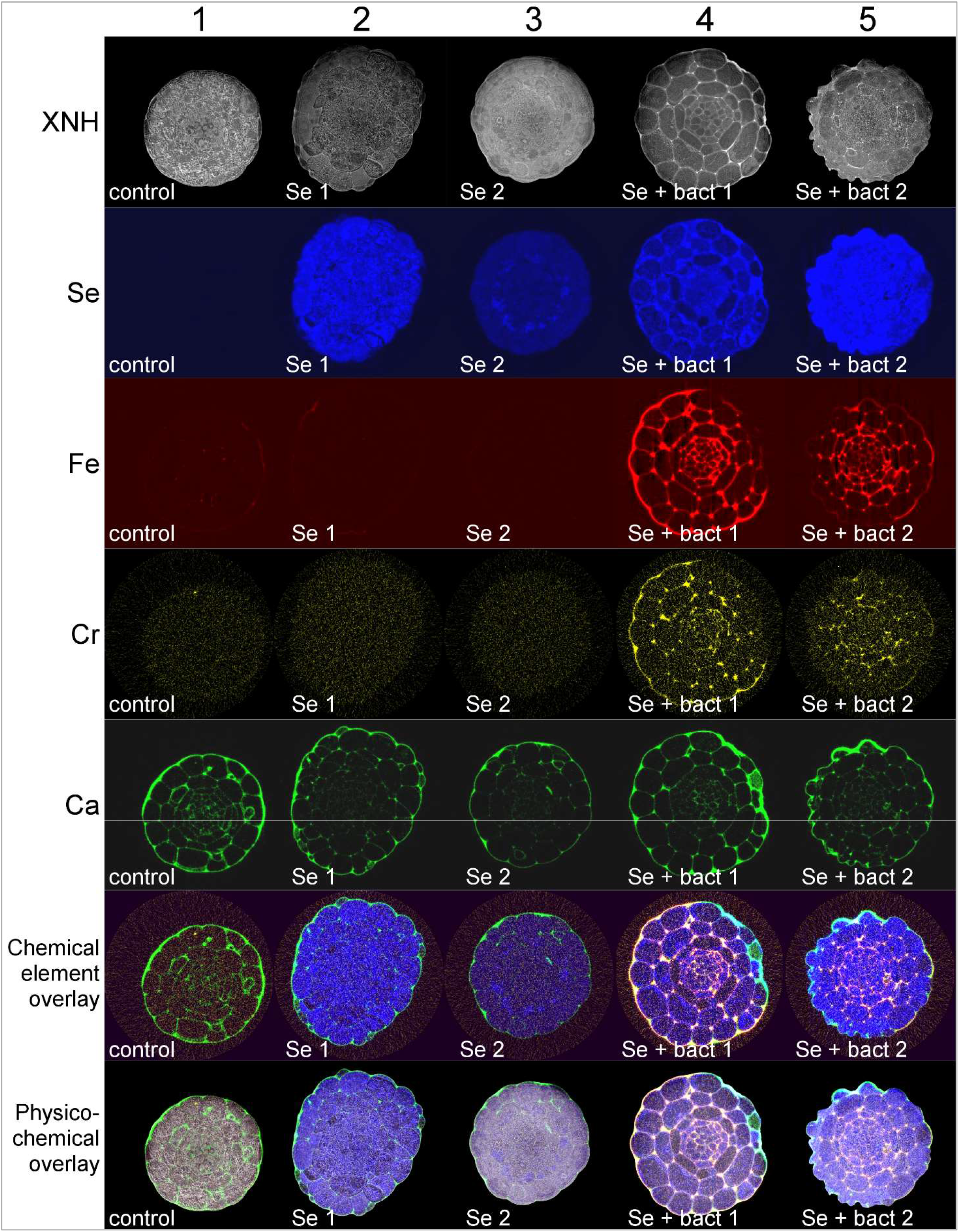
XNH electron density slice images and specific chemical element maps. This figure illustrates XNH slices of differently treated samples [1 (control), 2 & 3 (Se supplemented) and 4 & 5 (Se+bacteria supplemented)] and 2D chemical element maps of selected elements, Se, Fe, Cr and Ca at the exact position of XNH scans. Top row marked with XNH present physical electron density maps (XNH slices). Second row illustates XRNF signal of Se in samples. Third row shows signal of Fe, fourth row Cr and fourth row Ca signal in control roots. Composite overlay images of XRNF maps of Se, Fe, Cr and Ca are shown in the sixth panel. Composite images showning the physico-chemical maps are shown in bottom panel to illustrate the overall chemical composition differences between samples.

To better understand the element accumulation at subcellular level, close ups of Se and Fe maps were overlaid with electron density images (Figure 5). Again, control root (A) showed no sign of Se and low levels of Fe. In Se supplemented roots (B and C) Se appeared to be unevenly distributed, and Fe levels were barely detectable. In Se+bacterial supplemented roots (D and E) both Se and Fe signals were clearly visible. On subcellular level Se appeared visually to be primarily located in cytoplasm and internal membranes in various root cells. In bacteria co-cultured roots, apart from cytoplams and vacuoles, Se was also visually detected in some cell walls (Figure 5 D), supporting data presented in Graph 2. In all samples from which Fe could be detected, Fe signal was localized almost exclusively to cell walls and hyperaccumulated into the junctions of cells, showing a very different localization pattern than Se (Figure 5, Graphs 2 and 3). Following visual inspections, Se and Fe levels were quantified at different subcellular compartments from the XRNF maps (masks of the selected areas and measurements are shown in Graph 3). Results indicated that in all samples, cell walls & cytoplasmic areas contained on average equal levels of Se, which exceed Se levels in vacuolic compartments (Graph 3). Within vacuoles, the membranes contained more Se than fluid filled areas, at least in the case of Se+bacteria supplemented root #1 that exhibited clearest subcellular compartmentalization required in such fine detail measurements (Extended Figure 3). Segmentation analysis showed that in all measured samples Fe was predominantly localized to cell walls (Graph 3), supporting visual observations. When the Se and Fe levels of Se+bacteria supplemented roots were compared on across all tissues, a ~3.5-fold difference was observed in averaged mean values between Fe and Se. In cell walls specifically, the difference was even more pronounced as mean Fe levels were >12-times higher than Se levels (online material: Se_Fe statistics.xlsx). Levels of Cr and Br accumulation were also increased in cells of bacteria co-cultured roots – however the increase was not as high as for Fe. Cr appeared to be mainly localized to cell walls, while Br was more prominent within the cells (Extended figure 2). While the overall Ca signal intensity in cell walls appeared similar for all treatments (Figure 4, Extended Figure 2), few local higher Ca signal peaks were observed in cell walls of damaged epidermal cells of Se+bacteria supplemented roots (Extended Figure 4). The XNH 3D surface reconstruction of such damaged area is shown in Extended Figure 4. The concentration of Ca in these locations was ~2.5 times higher than in neighboring epidermal cell walls. It is not clear whether the damage was caused by the *Pseudomonas* bacteria visible on the surface of the root – or if it was caused by sample preparation and mounting onto cryo-sample holder.

**Figure 5:**
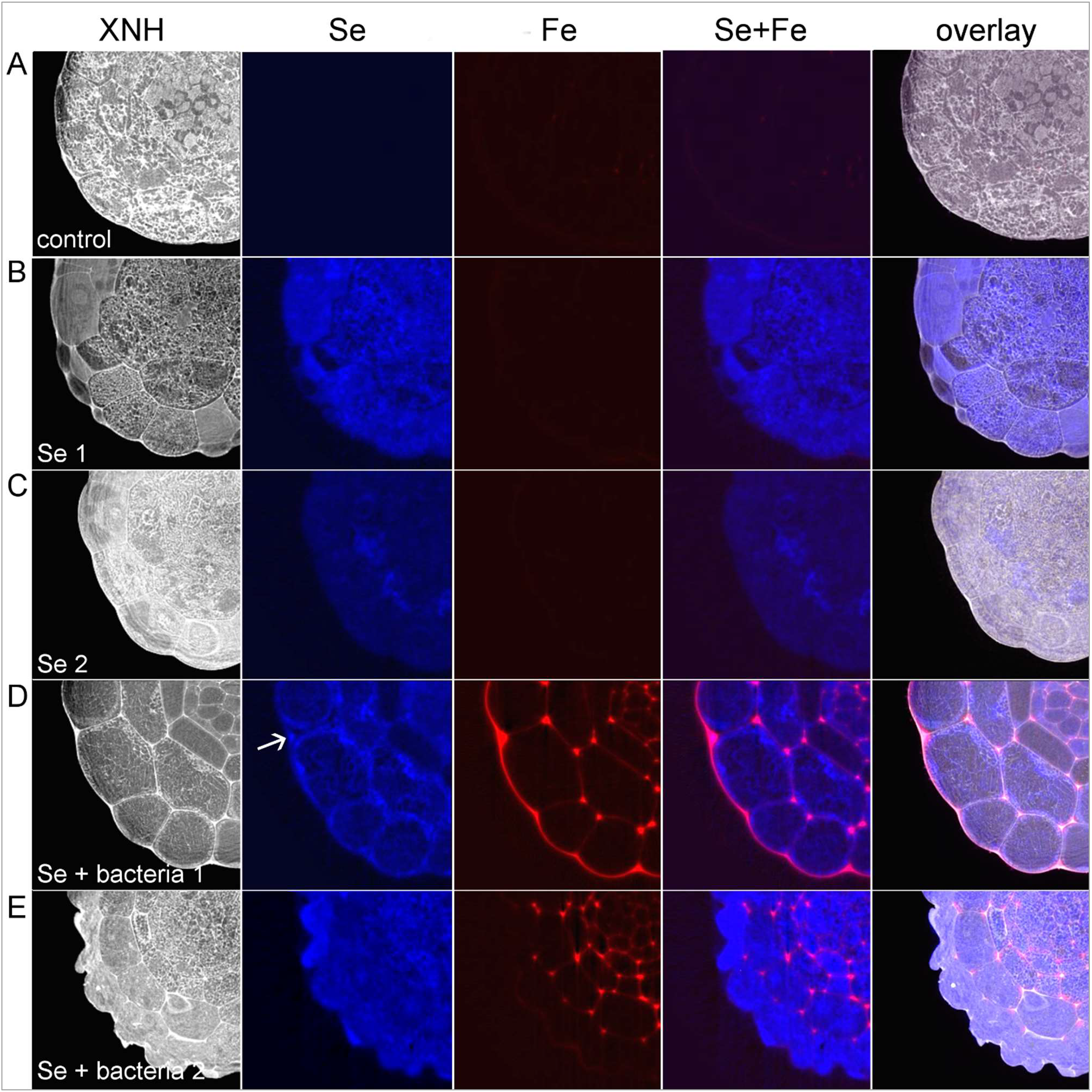
Closeups of Se and Fe in different treatments. First column on the left presents XNH imaged roots. Second column close ups of selenium and third column iron in root sections. In the fourth column Se and Fe signals are overlaid and presented in blue and red, respectively. The column in the right shows Se and Fe signals overlaid with XNH images for tissue visualization. Rows A – E represent imaged samples, control (A), Se supplemented (B and C) and Se+bacteria supplemented roots (D and E).

Finally, Se-Fe density scatter plots were generated from each sample (Extended Figure 5) and used to segment root areas based on their densities. When plotted between Se and Fe allocation domains, the plotted areas were in line with the XRNF maps (Figures 4 and 5). It was initially assumed that the dense intracellular aggregates located at vacuolic membranes and around nuclei of many samples, especially visible in Se+bacteria supplemented root (Extended figure 6), were Se enriched protein aggregates (as was suggested by Gupta & Gupta, 2016). However, the majority of these aggregates seen in both PXCT and XNH datasets did not match with XRNF Se maps, indicating that the aggregates consist of other chemical element(s). Interestingly, both Zn and Ca aggregates were found in the epidermal cells of most imaged samples (shown in Extended Figure 2), localizing mainly to vacuoles. In many cases these elements matched to the dense aggregates measured using XNH, indicating that these micronutrients can make dense aggregates in plant cells (as was suggested for Zn by Arrivault *et al.*, 2006)(data not shown).

## DISCUSSION

### Bacterial co-culturing affects root anatomy, tissue density and chemical composition

The electron density data was used to 1) visualize tissue and subcellular organelle anatomy in fine detail and 2) measure tissue densities within intact 3D samples. Each co-centric root tissue type (epidermis, cortex, endodermis, pericycle and stele, illustrated in Figure 2 and Extended Figure 7) could be identified based on their position, shape and cell numbers. Visualization of subcellular contents allowed comparisons between treatments in processes like cellular clearing and increase in vacuolic size and water content (seen especially endodermis, pericycle and stele in control roots), which occur during cellular differentiation, as the volume of cytoplasmic domain reduces upon cellular differentiation (Perrot-Rechenmann, 2010). For example, Se+bacteria supplemented roots exhibited altered cellular division and differentiation rates and higher density of vascular tissues (Figure 2) compared to both control and Se supplemented roots. Their increased intracellular density could possibly be attributed to 1) higher levels of sugars, fats and proteins in cytoplasm, 2) proportionally larger cytoplasmic cell fractions in cells, and to 3) increased number and size of subcellular aggregates (especially in epidermis and cortex cells). In addition to intracellular cytoplasmic density increase, Se+bacteria treated roots showed increased density of extracellular cell walls (table 1). This finding could at least partially attribute to the fact that the bacteria promote Fe fortification of cell walls. While some Se *did* localize to cell walls in bacteria supplemented roots, the difference in magnitude indicates that Se is not a major contributor to the observed cell wall density differences between different treatments (Fe levels in roots supplemented with Se+bacteria were >12-times higher than Se levels). Oppositely, as the electron densities of Se+bacteria treated root vacuolic membranes were ~17 higher than in control root, and Fe did not accumulate into vacuoles, it is quite possible that the observed increase in vacuolic membrane density in bacteria co-cultured roots was attributed primarily to Se aggregate formation.

### Se toxicity and bacteria elicitor promoted growth abnormalities

The observed treatment specific differences in root anatomy, including differences in root size (diameter), defects in maintaining proper cell divisions, and growth arrest, both in Se and Se+bacteria supplemented roots compared to control roots, could in part be attributed to biotic Se stress (toxicity) as the Se supplemented roots showed slight reduction in growth (Lusa *et al.*, 2019, Extended Figure 8). However, as the cell division defects, and meristem differentiation abnormalities were exaggerated in the Se+bacteria supplemented roots (Figure 3), it is unlikely that the phenotypes were caused by Se toxicity alone. While deviations from normal meristem patterning could at least partially explain the observed increased root hair phenotype seen in bacteria co-cultured *Arabidopsis thaliana* roots (Figure 3), there is an additional explanation to the observed pleiotropic root phenotype: the bacteria could possibly excrete plant phytohormone like elicitors to the growing media. Presence of bioactive auxin (or inactive precursors or auxin like compounds) could explain the observed phenotypes of our Se+bacteria supplemented roots: the Se+bacteria roots had larger meristems, exhibited extra root hair formation, showed root growth arrest (seen as white meristems, Extended Figure 8) and odd root branching architecture (Extended Figure 8). In fact, analyzed roots resembled closely several classical *Arabidopsis* auxin mutants, including abnormal root architecture of superroot (Boerjan *et al.*, 1995), *yucca*-mutants (Zhao *et al.*, 2001, Cheng *et al.*, 2006), and pleiotropic morphological and cellular differentiation defected *auxin resistant 1*-mutants (Lincoln et al.,1990). Moreover, the Se+bacteria supplemented roots showed an exposure time dependent increase in lateral root emergence phenotype (data not shown), similar to what was reported by Singh et al. in 2015. The bacterium induced auxin accumulation scenario is supported by the fact that several soil bacteria (similar to used *Pseudomonas* strain) are known on elicit auxin overproduction like root phenotypes in plants across the plant kingdom, most famous example being soil dwelling microbes belonging to the *Agrobacterium* genus (Esitken 1970, Cardarelli *et al.*, 1987, Amselem & Tepfer, 1992). The exact mechanism behind the putative auxin related phenotype remains to be determined, but could be due to auxin overproduction in plants (via bacteria originating lateral gene transfer promoted activation and/or overexpression of auxin biosynthesis genes) (Cardarelli *et al.*, 1987, Liu *et al.*, 2015) or caused by external auxin stimulus produced by the bacteria themselves. Overall, it can be concluded that the bacteria affected meristem growth (cell division and differentiation), vascular tissue density and subcellular composition (both intra and extracellular densities), but the exact mechanism remains unsolved.

### Superior properties of intact fresh tissues 3D maps

Previous efforts to study the amount and localization of Se in plant tissues have relied mostly on three very different approaches: μXRF (Wang *et al.*, 2013, Hu *et al.*, 2014, Zhao *et al.*, 2014) or spectroscopy imaging assays assays (Krajcarová *et al.,* 2017), cell organelle fractionation, and indirect gene expression assays (Gupta & Gupta, 2016) in response to exogenously applied chemical compounds. In fluorescence imaging assays conducted at room temperature, imaged samples were either rather large (whole roots, seedlings of leaves), fixed in resin, or freeze dried to minimize sample movement during imaging. All these approaches have their caveats: The spatial resolution in imaging of large samples will not be sufficient to reveal subcellular differences or fine features. Fixing a sample in resin includes dehydration steps, infusion with resins, and possibly sectioning – leading potentially to alteration of chemical composition and anatomical deformation. Same issues apply to freeze drying, in which all the water is often removed using *e.g.* ethanol series followed by lyophilisation. As water content affects critically the conformation, density, mechanical properties and chemical composition of biological tissues, every modification will inevitably contort the obtained data compared to samples imaged *in vivo*. While electron microscope imaging methods can provide direct information of chemical elements in high resolution, the samples are sectioned to thin slices. The imaged samples are also often dried to immobilize them, loosing information on water and soluble compounds. All these described procedures leave plenty of room for improvement on sample integrity but also on volume of information gained from such narrow “views” of 2D slices. Together, the previously used methods have provided mostly either low resolution, indirect or spatially limited evidence of both the physical and chemical properties of biological samples.

We overcame all these limitations by studying Se uptake and accumulation in fresh *Arabidopsis thaliana* root tissues using state-of-the-art synchrotron X-ray radiation based PXCT, XNH and XRNF tomographies at about 100 nm scale resolution. All imaging assays were conducted in cryogenic conditions in order to preserve sample features as intact as possible for reliable chemical element analysis. When samples are frozen, the movement caused by radiation damage, *i.e.* volatilization and reactive oxygen species formation due to high photon flux, is minimal allowing long measuring times. Long measuring times allow retrieval of > 1000 projections from each sample, leading to the desired image resolution of about 100 nm. As Se can be processed to volatile compounds, sample freezing also prevented enzymatic activity and putative loss of Se signal from samples. As a bonus, cryoimaging allowed inspection of transient chemical signaling events in plant tissues, preserving fresh samples features and local chemical element concentration peaks in samples as if the samples were in their native state.

### High-resolution imaging resolution reveals subcellular organelle features

The reconstructed high-resolution 3D electron density volumes from PXCT and XNH showed highly similar results on sample anatomy. This is noteworthy, as the electron density data was obtained using two different cryogenic X-ray imaging methods and the measurements were performed at different facilities from different biological sample batches. One difference was that unlike PXCT, XNH datasets appeared to highlight network like subcellular compartments (like ER and vacuolic membranes) based on their properties that enhance edges on compartments of different texture (e.g. fat vs. water). Due to the choice of large field of view, the XNH results were also of slightly lower resolution than the PXCT as evidenced by the subcellular features. The XNH datasets contained also some artefacts at samples edges (seen as “ripple effects” at the epidermal cell edges, Figure 2, panels D – F). These artefacts could be reduced by using higher incident X-ray energy.

The volumes allowed inspection of individual cells and subcellular compartments in fine detail. For example, phloem sieve elements could be identified (Figure 2 G and Extended Figure 6) based on the cell wall properties and developmental status of different cells in the stele. In *Arabidopsis thaliana* root, phloem sieve element cells are first vascular cells to differentiate at the transition zone (at the exact position where the in dataset were collected). Because of this, phloem sieve element cells have denser cell walls than the adjacent phloem companion cells, procambial or phloem pole pericycle cells (visible in Extended Figure 6, cells with asterisks). Sieve element differentiation is accompanied by clearing of cellular contents in a process called enucleation and phloem cell clearing (Furuta *et al.*, 2014, Blob *et al.*, 2018) (visible in Figure 2 G, cell type 5). The visualization of phloem cell clearing process was originally reported using laborious high-resolution cryo-EM section assay that relies on imaging 2D slices of electron microscopy samples (Furuta *et al.*, 2014,). Remarkably, we were able to obtain information on phloem sieve element cell wall density properties and visualize cellular clearing from 3D stacks of intact fresh samples (albeit not at same ultra-high resolution as with 2D EM imaging). In addition to cell wall density properties, very fine subcellular anatomical features were distinguishable within the 3D datasets of soft plant tissues simply based on minor differences in their electron densities. With the obtained resolution, it was possible to produce strikingly detailed images (like Figure 2 H – L) and distinguish various subcellular organelles within imaged intact samples, including mitochondria, vacuoles, and different nuclear compartments (i.e. nuclear envelope, ribosomal protein enriched nucleolus, nucleoplasm, nuclear cavities, and chromatin). To the authors’ knowledge, similar high-quality absolute electron density 3D data has not been reported from fresh multicellular plant samples, especially in such detail. **This makes our PXCT and XNH results truly novel in two ways: 1) Subcellular features previously only recorded with cryo-EM methods can now be determined in 3D from intact fresh plant tissues. 2) Our datasets report actual sample electron densities instead of mere anatomical features.**

### Elemental information at subcellular organelle resolution

Our multi-modal 3D cryo-imaging approach yielded high-resolution electron density and elemental maps of intact fresh tissues at cell organelle resolution providing direct evidence of e.g. aggregate composition. Due to time constraints XRNF tomographic imaging was done using a larger pixel size of 200 nm and only a single or a few slices were imaged. To visualize and quantify distribution of Se and other micronutrients in root tissue and subcellular compartments, the higher resolution anatomical XNH electron density features were matched to individual chemical elements (mg/cm^3^ values) obtained via XRNF measurements.

We observed differences in Se levels between samples and treatments, but overall Se accumulation profiles were similar between different tissues (epidermis, cortex, endodermis, and stele) (Graph 1). Se accumulation appeared to be enhanced in roots co-cultured with bacteria. Thus, based on our data, we could conclude that bacterial co-culturing does indeed promote Se accumulation to root tissues. According to previous reports plants accumulate Se in vacuoles (Gupta & Gupta, 2016). Using direct fluorescence measurements, we were able to quantify Se levels from various subcellular compartments (cytoplasm, cell walls and vacuoles). In both Se and Se+bacteria co-cultured roots highest values were measured from cytoplasm and cell walls, not from vacuoles. It should however be noted that due the cell wall masks used in segmentation analysis, some Se signal from cytoplasmic was likely included in the cell wall fraction, emphasizing subcellular cell wall measurements over other fractions. In addition, due to limited resolution, vacuoles were pooled as individual areas (including both membranes and vacuolic fluid). When analysis was done from individual hand selected points across the root, Se measurements looked slightly different: highest Se values came from cytoplasm, followed by aggregates in vacuolic membranes (lowest values were measured from vacuolic fluid). Nonetheless, we confirmed that Se does accumulate and aggregate in plants, however not exactly in similar manner as bacterial and mammalian cells.

In addition to differences in Se accumulation and subcellular localization, we observed changes in other micronutrient levels in response Se+bacteria supplementations, including high concentration Fe deposits in cell walls and Zn and Ca aggregates (also clearly visible in the videos of 3D XRNF volumes provided in online material). Interestingly, unlike initially expected, the observed high electron density aggregates seen in PXCT and XNH datasets did not match with XRNF Se maps but rather to Zn and Ca measurements. It seemed like the overall number and size of such dense aggregates was larger in bacteria co-cultured roots than in control or Se alone treated roots (Figure 2), indicating that bacteria affect root chemical composition in a complex manner. While bacteria have been shown to change the redox status of Se in the growing medium (Lusa *et al.,* 2017), possibly affecting uptake of Se, the methods we used in our research could not determine chemical speciation of the imaged elements. For such data chemical element speciation sensitive methods should be applied. Therefore, the questions about the possible influence of bacteria supplementation to the speciation of elements in the roots and the effect of chemical speciation to uptake of Se and Fe (and other elements) into tissues remains to be addressed.

### Catching mechano-sensing “mid-flight”

One remarkable finding was that cryo-imaging allowed visualization of transient “moments frozen in time” preserved over extended imaging times. In the imaged samples, majority of Ca signal was seen evenly distributed at cell membranes/cell walls (Figure 4, Extended Figure 7). This is interesting, as Ca is known to act as a signalling component for mechanical stress, free oxygen radicals, and gravistimulus in plant cells, and Ca is locally released from cell walls and subcellular compartments rapidly upon such stimuli (Demidchik *et al.*, 2003, Monshause at al., 2011, Kurusu *et al.*, 2013). Our measurements show high local Ca peaks in Se+bacteria supplemented roots in cells that were slightly damaged just prior to freezing (Extended Figure 7). In these cells, Ca levels were 2.5-fold higher than in the surrounding intact cell walls (Extended Figure 7, graph). While wounding and subsequent Ca burst was most likely induced by direct mechanical stimulus during sample preparation, we cannot exclude the possibility that the bacteria residing at the surface of the root might have contributed to cellular damage. Regardless of the cause, this example illustrates that non-invasive cryogenic imaging is especially well suited for preserving *in vivo* features of fresh tissues over extended imaging times (in our case up to 18h). In addition, it is worth noting that these same cells could be segmented from the voxel-wise Se vs. Fe concentration scatter plots as they form a clear outlier region distinct from other regions (Extended Figure 9). This indicates that utilization of multichannel data and advanced segmentation algorithms has potential to classify and reveal anatomical and physiological information which could be otherwise difficult to see. However, such considerations are beyond the scope of the current work.

### Bioremediation and future of micronutrient accumulation assays

Using a multimodal cryogenic 3D and 2D imaging approach we were able to obtain comprehensive data of fresh *Arabidopsis thaliana* root samples in their native state with higher spatial resolution than what is currently reported in literature. Our observations on the chemical landscapes of individual cells and cellular organelles can promote understanding of metabolic processes involved in storing, processing and translocating Se and other micronutrients within the plant body. In addition, our data offers possible explanations to how soil bacteria affect root anatomy and growth. We suspect that the bacteria elicit phytohormone like growth regulators that lead to observed cellular division and tissue patterning defects. These abnormal cellular divisions then lead to super numerous root hair formation promoting micronutrient uptake (explaining the observed increase in Se and Fe accumulation). It is also possible that the bacteria promoted increase in micronutrient accumulation exceeds the toxicity limit for Se (and possibly Fe), leading to the observed growth arrest/tissue deformation and micronutrient hyperaccumulation. Regardless of the exact mechanisms which require more work until solved, the observed effects of bacterial supplementation open up an important promising possibility: *Pseudomonas* sp. strain T5-6-I soil bacteria could be utilized in both Se bio-remediation and Fe bio-fortification (likely even without Se supplementations), improving the nutritional values in agriculturally important crops grown on Fe poor areas. **Our findings demonstrate the great potential of soil-microbe co-culturing on bioremediation & biofortification efforts and illustrate how such interactions and their effects can be studied *in planta***. This in turn will improve our agricultural practices, especially in eroded areas where soil health, micro- and macronutrient availability and plant uptake would benefit tremendously from the introduction of beneficial soil microbes.

## METHODS MATERIAL

Surface sterilized (0.1 % Triton X in 95% ethanol) *Arabidopsis thaliana* wild type (Col-0 ecotype) seeds were sown on 1xMS-agar (Murashige and Skoog Basal Salt Mixture, Sigma M5524) (Xi et al. 2016) plates (50 mL) containing 1% sucrose (w/v) and 0.7% (w/v) agar (Sigma A1296).The seeds were kept at 4 °C in darkness for minimum of 2 days (stratification) and then transferred to a growth chamber set at 23°C in 16/8 h light and dark cycles (long day cycle, LD). Selenium and Bacterial suspensions of *Pseudomonas* sp. strain T5-6-I (5·10^7^ CFU of exponential growth phase bacteria) were added on one-week old seedlings, by spreading in total 5 mL of the suspensions on the surface of the plant agar. The plants were thereafter co-cultured with Se and bacteria for 7 days before imaging (Extended figure 1). The plants were transported to synchrotron facilities in hand luggage and kept under LD during sample preparation and imaging procedures. As we have reported before (Lusa *et al.,* 2019) the treatments affect root phenotypes (Extended figure 1). Therefore, the effect of each treatment was verified on site with light microscopy. An overall visual inspection of plants used for PXCT and XNH verified that soil bacteria co-culturing affects root anatomy and architecture (Extended Figure 2). While plants grown on normal media showed normal primary and lateral root elongation rates and lateral root branching pattern (Extended Figure 2 A, D), plants supplemented with Se showed slight reduction in root growth rates (Extended Figure 2 B and E). However, the overall lateral root branching pattern seemed unaltered in Se supplemented plants. In clear contrast, the plants supplemented with Se+bacteria showed a reduction in root growth rate and abnormal lateral root branching phenotype at a high frequency (not quantified) (Extended Figure 2 C, G and H). Bacteria supplemented roots also exhibited a hairy root phenotype, as was reported with kale (Lusa *et al.,* 2019) (Extended Figure 3, panels G - L). Root hair imaging was done at the Light Microscopy unit at the Institute of Biotechnology, Helsinki, Finland. Imaged roots were extracted from media and stained with propidium iodine (PI) before imaging with a Leica SP5 inverted confocal microscope using a solid-state blue laser for GFP (480 nm/270mW).

In addition to increased root hair formation, bacteria co-cultured roots showed high rate of lateral root meristem arrest (white root tips, Extended Figure 2 G) (not quantified). Such arrested roots were not included in assays and imaging was only done with healthy looking roots of similar age (Extended Figure 2 D - F) at around 2 d post emergence. Healthy looking lateral root tips were mounted on cryogenic sample holders (OMNY-pins [Holler *et al.,* 2017] and aluminium Huber-tubes at the Swiss Light Source (SLS) and at the European Synchrotron Radiation Facility (ESRF), respectively, and snap frozen in liquid nitrogen.

## IMAGING AND DATA RECONSTRUCTION

All synchrotron imaging assays (PXCT, XNH, and XRNF) were done at a position corresponding to the transition zone (blue boxes in figure 1), which acts as a divider between meristematic cells and elongating and differentiating cells. This zone is especially interesting as cells of this zone act as sensors for various environmental stimuli, they are active in cytoskeletal rearrangements and mediate growth responses leading to initiation of both root hairs and organs (Baluška and Mancuso, 2013).

The cryo-PXCT measurements were performed at the cSAXS beamline at the SLS at the Paul Scherrer Institut (PSI), Switzerland, using the OMNY instrument (Holler *et al.,* 2018). Samples were glued to OMNY pins (Holler *et al.,* 2017) at room temperature and then frozen in liquid nitrogen. The pins were transferred to the vacuum chamber and measured at a sample stage temperature of 90 K. Cryogenic conditions can help maintain tissue and cell shape, and reduce possible volatilization of Se. Samples were centered at about 100 μm distance from the root tips (variation in root tilt affected distance). The heights of imaged areas ranged between 15-25 μm, depending on the angle and/or width of the roots. A gold Fresnel zone plate fabricated at the Laboratory for Micro- and Nanotechnology at the PSI with a diameter of 250 μm and 50 nm outermost zone width was coherently illuminated at 6.2 keV photon energy. Spatial horizontal coherence was achieved by a horizontal upstream slit and temporal coherence by a Si(111) double crystal monochromator. For the measurement the samples were placed 1.6 mm after the focal spot of the Fresnel zone plate, where the beam size had reached 6.5 μm diameter. The diffracted X-rays have been recorded by an in-vacuum Eiger detector (Guizar-Sicairos *et al.,* 2014) placed at a distance of 5.213 m after the sample.

Ptychographic projections were recorded scanning the sample in the beam following the Fermat spiral trajectory (Huang *et al.,* 2014) with a step of 2 μm. The field of view and number of projections was adjusted according to sample diameter and available measurement time. Table 1 gives an overview of the different samples measured and the parameters used. In order to keep imaging times reasonable (limiting to max 23-24 hours per sample, corresponding to ~1200 projections), the resolution vs stack height was chosen separately for each sample. The projections were reconstructed using between 1000 and 1600 iterations of the difference map algorithm (Thibault *et al.,* 2008) followed by 600 iterations of maximum likelihood refinement (Thibault & Guizar-Sicairos, 2012). For the reconstructions, either 500 x 500 or 400 x 400 pixels of the detector were used, resulting in a reconstructed pixel size of 27.8 or 34.7 nm, respectively. The projections were aligned (Guizar-Sicairos *et al.,* 2011 and 2015) and combined to 3D volumes using a modified filtered back projection (Guizar-Sicairos *et al.,* 2011) using a GPU-based matlab routine as described in Ref. (Odstrcil *et al.,* manuscript in preparation). The 3D resolution was estimated for each tomogram using Fourier shell correlation (FSC) between two subtomograms with half the number of projections each and using the half-bit criterion (van Heel & Schatz, 2005). An illustration of raw data and tissue identification from control root samples is shown in Extended Figure 7. The results of the FSC analysis are also given in table 2 (Extended material).

XNH and XRNF were performed under cryogenic conditions (LN2 temperature) under ultra-high vacuum (Figure 1, B) at ESRF ID16A beamline. XNH volumes measured were centered at 115μm distance from the root tips (full volumes shown in Figure 3). A brilliant beam (10^11^ phot/s) of highly focused X-rays (30 (H) x 30 (V) nm^2^) at 17 keV energy was produced by an undulator source and a KB mirror optic (cf. Figure 1 B) (Morawe *et al.*, 2015). The focus was used as a virtual point source for the XNH imaging by placing the sample some distance downstream from the focus and recording a transmission image on the detector. These XNH images show intricate interference patterns that carry information about the X-ray phase shift happening in the sample. A tomographic scan was performed over 180-degree range with 0.1-degree step size. At each rotation position the sample was randomly (but controllably) displaced to reduce systematic artefacts from the incident beam and detector non-uniformities. Such tomographic scan was performed at 4 different propagation distances by varying the sample distance from the X-ray focus. A linear phase retrieval algorithm (Paganin et al. 2002) extended to multiple propagation distances and followed by iterative corrections was then applied to the set of 4 projection images at each rotation angle. This produced 1800 quantitative phase projection images covering the 180-degree rotation, which were subsequently used for tomographic reconstruction with PyHST software package (Mirone *et al.*, 2014) to get a 3D volumetric image of the electron density in the sample. The resulting voxel size of the images was 50 nm, which was chosen so that the whole cross section of the sample would fit into the image, resulting in a total imaged volume of about 100 ×100 ×100 μm^3^.

The XRNF measurements were conducted by placing the sample in the X-ray focus, and then raster scanning the sample to produce 2D maps and sinograms. The raster scanning was performed with 200 nm spatial step size, 27 ms dwell time, and 0.72-degree step over 180 degrees rotation. Either 1 or 7 slices (spaced by 200 nm) were recorded in these scans depending on the sample. The corresponding XRNF 2D slices and 3D stacks were located at the center of the XNH field of view, at 115 μm distance from the root tips. The incident energy of the X-rays was high enough to ionize all element species of interest in the sample. A set of energy discriminating detectors placed at the sides of the sample were used to record the fluorescence spectra coming from the sample. The summed spectra from these detectors was then used to do peak fitting using the PyMCA software package (Solé *et al.*, 2007). The measurements were brought to quantitative units by using a previously measured calibration sample. The end results of this peak fitting step were therefore images of different elements in units of surface density (ng/mm^3^). The fitted images were then organized as sonograms, which were then reconstructed using the PyHST software package with an iterative algorithm that minimizes the total variation in the object by using Chambolle-Pock type method (Chambolle & Pock, 2010). This resulted in cross sectional slice images of local concentration (gm/cm^3^) for each of the analyzed elements. Due to the high incident flux and small background signal, the XRNF images have a final detection limit of about 1 ppm for each element.

## DATA ANALYSIS, VISUALIZATION AND SEGMENTATION

For electron density analysis PXCT datasets were imported to and calibrated in Fiji using experiment specific values for each dataset. These calibrated volumes were then used for electron density measurements of various tissues and subcellular organelles from each sample. In case of subcellular organelle measurements average values (min, max, median and mean) were calculated from over 30 measuring individual areas around the sample and presented as averages (table 2) and as bar charts including error bars (Extended Figure 4). The sizes of measure areas ranged from tens of pixels (cell walls, vacuolic membranes) to larger areas covering several thousand pixels per measured point (cytoplasmic and vacuolic fluid), depending on the size of analyzed cellular organ.

XNH volumes and XRNF slices were imported, analyzed and processed in Fiji for electron density visualizations and chemical element measurements. Chemical element measurements done both at root, tissue and cell organelle level (for Fe and Se,) were done in Fiji. Tissue averages and subcellular measurement averages are presented in Graphs 1 – 3 created using Microsoft Excel. Se levels were measured from each tissue type from in all samples, from 20-30 areas per each cell type (different cell types are areas from which measurements were taken are illustrated in Extended Figure 7). Subcellular Se measurements were done from Se+bacteria sample with the most distinct anatomy for reliable scoring of subcellular compartments. In this case, numerous small areas around the root were selected (Extended Figure 8) and measured as pooled areas from epidermis, cortex and endodermis. Resulting measurements for vacuole membranes, vacuolic fluid and cytoplasm are presented in Graph 2. Fe levels were measured from 5+5 points around the root per sample for cell walls and intracellular areas (Graph 3). All graphs and table 2 were generated from measurements done in Fiji by exporting measured values to Microsoft Excel.

For visualizations individual images were exported in 16-bit tiff format and processed in Corel PaintShop Pro X9 to create multilayered composite and panel images. Pseudo-colored images of root cell types shown in Figure 1 were created from reconstructed PXCT electron density image slices by hand using Corel PaintShop Pro X9. Each cell type was colored onto a separate layer on top of the underlaying electron density image. Pseudo-colored slice images (Figure 2, panels G and H, Extended Figure 3) consisting originally of a single channel (grey) were produced with Fiji color palette function reflecting the differences in intensity of the object. 3D visualizations of XNH datasets (of which slice images are shown in Figure 3) were generated by exporting image stacks of calibrated datasets from Fiji as tiff. and bmp. stacks and processed with CTvox from Bruker.

Concentrations of Se and Fe in various tissues and subcellular compartments in Graphs 1, 2 and 3, and online material Se_Fe statistics.xlsx were determined by tracing the anatomical features from XNH slices using a combination of pixel and vector graphics programs (Corel PaintShop Pro X9, Gimp2, Inkscape) and overlaying the traced image with the corresponding XRNF slice to produce bitmap masks for voxel discrimination. The distributions of concentrations in voxels over masked areas were calculated using custom Python 3 scripts utilizing Numpy, Scipy, and Pandas packages, and the violin plots were produced using Matplotlib and modified Seaborn packages.

For Se vs. Fe scatter plots, region of interest selections and the corresponding voxel visualizations (Extended Figure 9), the corresponding XNH and XRNF slices were aligned by forward projecting the reconstructed slices in x- and y-directions and finding the translational offset in x and y by cross-correlating the resulting 1D projections. The contribution of background signal was minimized by calculating masks over the reconstructed root by thresholding the data and calculating a convex hull around the remaining voxels. For scatter plots 2D histograms were evaluated from the voxelwise Se-Fe concentrations. To visualize the locations of voxels in the sample on the basis of scatter plots, regions of interest were defined as polygons in Se-Fe plots and XRNF voxels were masked on top of the anatomical XNH slice based on their inclusion in the polygons. All the computational and visualization were made using custom Python 3 scripts utilizing Numpy, Scipy, Matplotlib, Shapely, and Scikit-image packages. The analysis codes are available from the authors by request.

## Supporting information

Extended Figures for manuscript

extended figure 3 data - selenium in subcellular compartments

extended figure 4 - Ca levels in root cells

graphs 1, 2 and 3 statistics

control root fluotomo

Se root fluotomo

Se+bact root fluotomo

control root cross section, density map

control root lateral section, density map

Se root cross section, density map

Se root lateral section, density map

Se + bact root cross section, density map

Se + bact root lateral section, density map

control root image 1

control root image 2

Se root 1 image 1

Se root 2 image 1

Se root 2 image 2

Se root 3 image 1

Se + bact root 1 image 1

Se + bact root 1 image 2

Se + bact root 2 image 1

Se + bact root 2 image 2

Se + bact root 2 image 3

Se + bact root 3 image 1

Se + bact root 3 image 2

Se + bact root 3 image 3

sample identities

---

**Graph 1:**
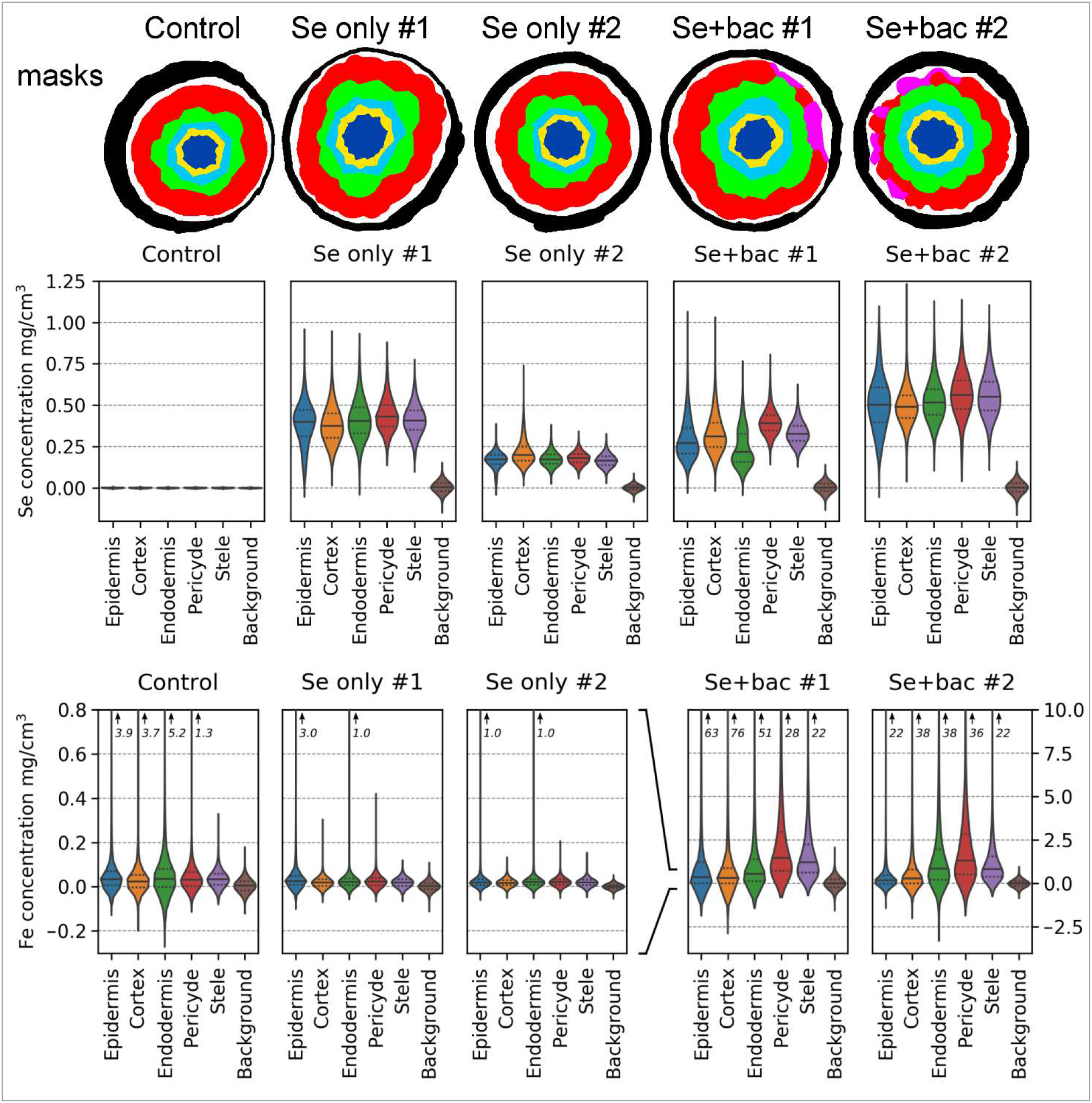
Selenium and iron contents in various root tissues of different treatments. Top: Tissues from which Se and Fe levels were measured: epidermis (red), cortex (green), endodermis (light blue), pericycle (yellow), stele (dark blue) and background (black). Pink areas represent damaged cells that were excluded from analysis. Middle: The solid horizontal lines within the violin plots of various tissues and adjacent cell walls represent medians, and the dashed lines mark 25 and 75 percentile tails. Peak values are indicated with numbers. The bottom and top values represent minimum and maximun signal intensities, respectively (anything below 0 should be considered a measurement artefact). Bottom: The Fe levels of control and Se supplemented roots are in similar range, with slightly higher values in control root. The Se+bacteria supplemented roots show considerably higher levels of Fe (notice the different scale on y-axes).

**Graph 2:**
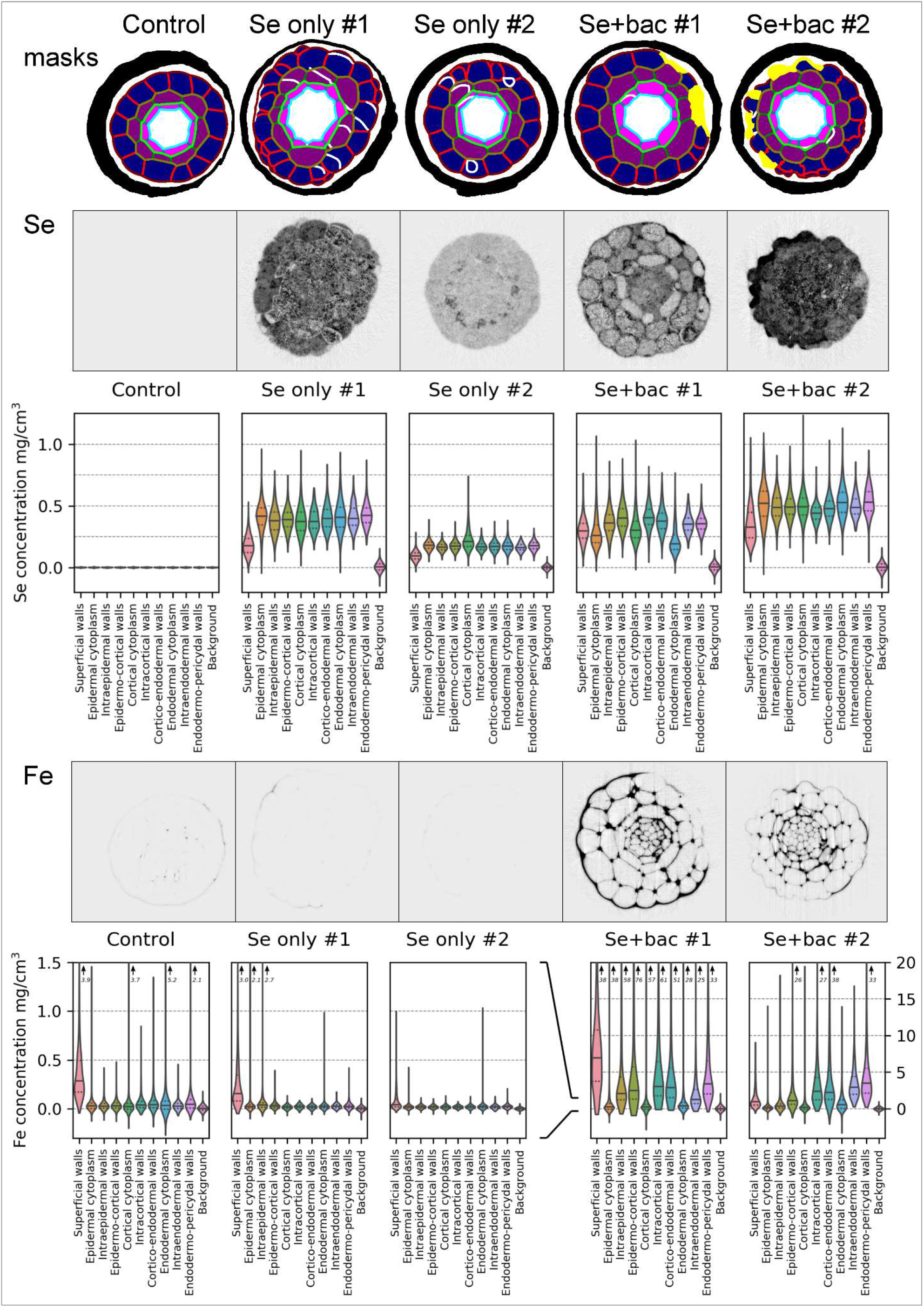
Selenium and iron contents measured from cells and walls from various tissues from different treatments. Se and Fe levels were measured from within all ground tissue layers, and from different cell wall positions (e.g. cell walls connecting only epidermis cells, and separately from cell walls connecting epidermis and cortex cells etc.). The masks representing measured areas are presented on the top: superficial cell walls (burgundy), epidermal cell contents (dark blue), epidermal cell walls (red), epidermocortical cell walls (brown), cortical cell contents (dark purple), cortical cell walls (dark green), corticoendodermal cell walls (bright green), endodermal cell contents (fuchsia), endodermal cell walls (turquoise) and endodermo-pericydal cell walls (light blue). Yellow areas are damaged cells, and white cell walls that were both excluded from measurements. Black areas, background. The solid horizontal lines within the violin plots of various tissues and adjacent cell walls represent medians, and the dashed lines mark 25 and 75 percentile tails. The bottom and top values represent minimum and maximum signal intensities, respectively (anything below 0 should be considered a measurement artefact). Medians are shown in solid lines and 25 and 75 percentile tails are separated with dashed lines. Peak values are indicated with numbers. Accumulation of Se appears quite even between intracellular and cell wall areas in Se and Se+bacteria supplemented roots. Opposite to Se, Fe appears to accumulate predominantly into cell walls, and in all samples, the superficial cell walls facing the growing medium showed highest values for all treatments. Fe accumulation was extremely pronounced in Se+bacteria supplemented root samples, where the median (solid lines) and highest values (peaks) were ~10X higher than in control root (notice the differentially scaled y-axes).

**Graph 3:**
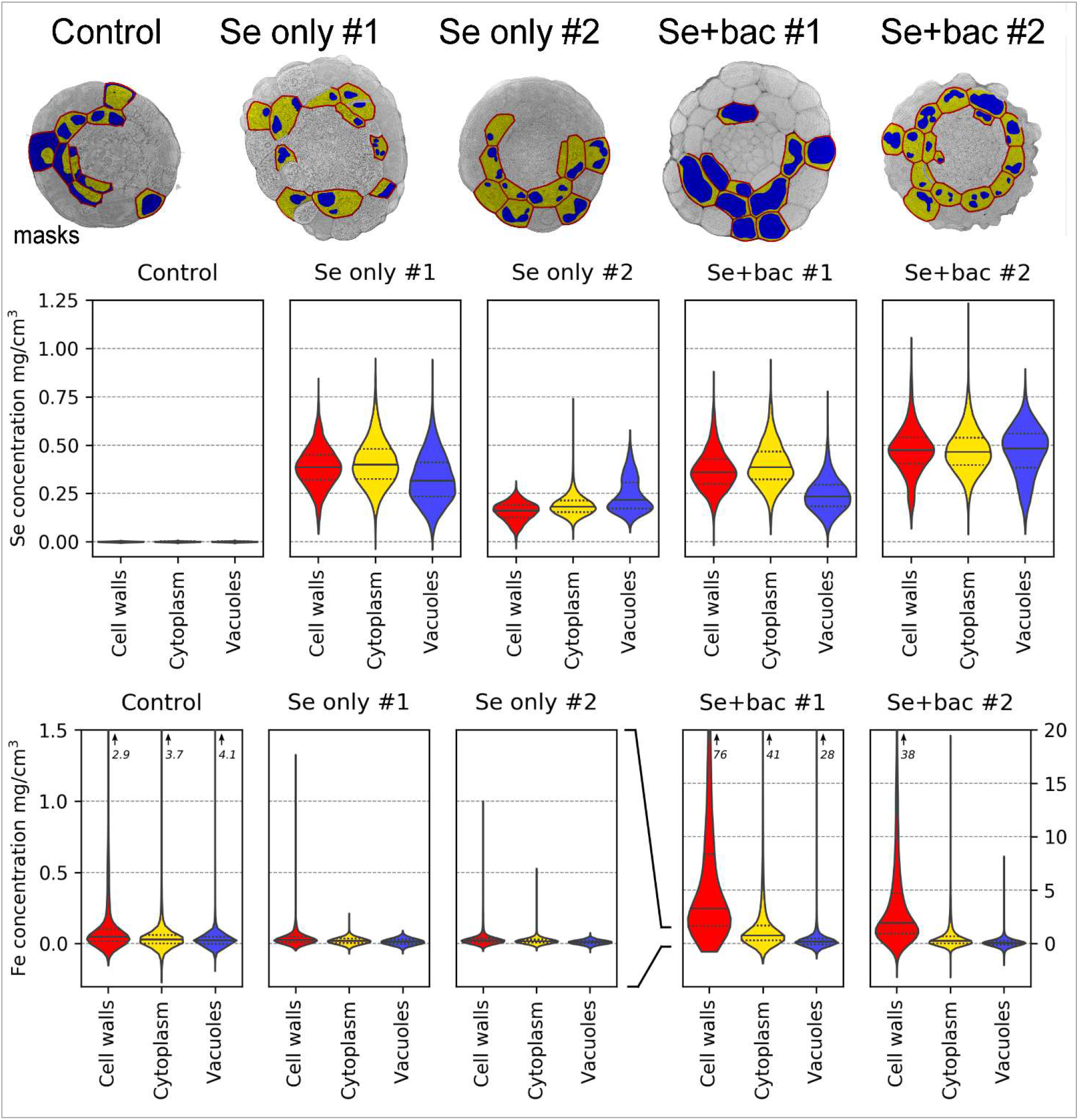
Selenium and iron contents in different subcellular organelles. Se and Fe levels were quantified at cell organelle level from all XRNF imaged samples. Masks used in quantification are shown on top row. Not all cells would be included into analysis due to suboptimal tissue resolution. Cell walls are shown in red, cytoplasmic areas in yellow and vacuolic compartments in blue. The solid horisontal lines within the violin plots of various tissues and adjacent cell walls represent medians, and the dashed lines mark 25 and 75 percentile tails. The bottom and top values represent minimum and maximun signal intensities, respectively (anything below 0 should be considered a measurement artefact). Based on these segmentation analyses, cell walls and cytoplasm contained on average more Se than vacuolic compartments. Notice the different scale on y-axes.

## ACKNOWLEDGEMENTS

Hanna Help and Henrik Mäkinen were supported by the Academy of Finland (Grant 1295696). Merja Lusa supported by the project “Biosorption and bioaccumulation of heavy metals and radionuclides - use of environmental bacteria in the bioremediation of mining waste-waters” from KAUTE foundation. Ari-Pekka Honkanen was funded by University of Helsinki Doctoral Program in Materials Research and Nanosciences (MATRENA).

## AUTHOR CONTRIBUTIONS

**H.H.** designed the experiments, participated in material preparations, conducted all sample preparations, participated in measurements, performed data analysis, and wrote the manuscript. **M.L.** participated in experimental design, prepared the plant material, participated in measurements and manuscript preparation. **A.-P.H.** participated in measurements, cryo-PXCT data reconstruction, analysis of cryo-PXCT, -XNH and -XRNF data, and manuscript preparation. **A.D**. and **M.H.** performed the cryo-PXCT measurements and data reconstructions and participated in manuscript preparation. **H.M.** participated in cryo-PXCT measurements and data reconstructions. **M.S.** and **P.C**. performed the cryo-XNH and -XRNF measurements and data reconstructions. **S.H.** participated in experimental design and manuscript preparation as well as in overall project supervision. **H.S.** participated in experimental design, cryo-XNH and -XRNF measurements and data reconstructions, and manuscript preparation.

## COMPETING INTERESTS

The authors declare that there is no conflict of interest regarding the publication of this article. The also authors declare that they have no known competing financial interests or personal relationships that could have appeared to influence the work reported in this paper.

## MATERIALS & CORRESPONDENCE

All correspondence regarding the manuscript and scientific content should be addressed to Dr Hanna Help. Requests for materials should be addressed to Dr Merja Lusa. Requests for segmentation code should be addressed to Ari-Pekka Honkanen. All data requests should be addressed to Dr Heikki Suhonen.

## Notes

#### Summary of Updates

First author affiliation

## REFERENCES

1. Amselem J & Tepfer M. (1992). Molecular basis for novel root phenotypes induced by Agrobacterium rhizogenes A4 on cucumber. Plant Molecular Biology, 19 (3); 421–432.

2. Arrivault S, Senger T & Krämer U. (2006). The Arabidopsis metal tolerance protein AtMTP3 maintains metal homeostasis by mediating Zn exclusion from the shoot under Fe deficiency and Zn oversupply. The Plant Journal, doi.org/10.1111/j.1365-313X.2006.02746.x

3. Baluška F and Mancuso S. (2013). Root Apex Transition Zone As Oscillatory Zone. Frontiers in Plant Science. 2013; 4: 354. doi: 10.3389/fpls.2013.00354

4. Blob B, Heo J & Helariutta Y. (2018). Phloem differentiation: an integrative model for cell specification. Journal of Plant Research, 131(1): 31–36. doi: 10.1007/s10265-017-0999-0

5. Bodnar M, Konieczka P & Namiesnik J. (2017). The Properties, Functions, and Use of Selenium Compounds in Living Organisms. Journal of Environmental Science and Health Part C Environmental Carcinogenesis & Ecotoxicology Reviews. 2012;30(3):225–52. doi: 10.1080/10590501.2012.705164.

6. Boerjan W, Cervera M, Delarue M, Beeckman T, Dewitte W, Bellini C, Caboche M, Van Onckelen H, Van Montagu M, Inzé D. (1995). Superroot, a recessive mutation in Arabidopsis, confers auxin overproduction. The Plant Cell, 7: 1405–1419. doi.org/10.1105/tpc.7.9.1405

7. Cardarelli M, Mariotti D, Pomponi M, Spanò L, Capone I & Costantino P. (1987). Agrobacterium rhizogenes T-DNA genes capable of inducing hairy root phenotype Molecular Genetics and Genomics, 209(3):475–80.

8. Chen S, Deng J, Yuan Y, Flachenecker C, Mak R, Hornberger B, Jin Q, Shu D, Lai B, Maser J, Roehrig C, Paunesku T, Gleber S, Vine D, Finney L, VonOsinski J, Bolbat M, Spink I, Chen Z, Steele J, Trapp D, Irwin J, Feser M, Snyder E, Brister K, Jacobsen C, Woloschak G and Vogt S. (2014). The Bionanoprobe: hard X-ray fluorescence nanoprobe with cryogenic capabilities. JOURNAL OF SYNCHROTRON RADIATION ISSN: 1600-5775 21(19): 66–75. doi.org/10.1107/S1600577513029676

9. Cheng Y, Dai X & Zhao Y. (2006) Auxin biosynthesis by the YUCCA flavin monooxygenases controls the formation of floral organs and vascular tissues in Arabidopsis. Genes Development. 2006 Jul 1;20(13):1790–1799. doi: 10.1101/gad.1415106

10. Chambolle A & Pock T. (2010). A first-order primal-dual algorithm for convex problems with applications to imaging. Journal of Mathematical Imaging and Vision. 40(1): 120–145

11. Cruz E, Becker S, Becker S & Sussulini A. (2018). Imaging of Selenium by Laser Ablation Inductively Coupled Plasma Mass Spectrometry (LA-ICP-MS) in 2-D Electrophoresis Gels and Biological Tissues. Methods in Molecular Biology. 1661:219–227. doi: 10.1007/978-1-4939-7258-6_16.

12. Cui J, Liu T, Li Y & Li F. (2018). Selenium reduces cadmium uptake into rice suspension cells by regulating the expression of lignin synthesis and cadmium-related genes. Science of the Total Environment, 644: 602–610.

13. da Silva J C, Pacureanu A, Yang Y, Bohic S, Morawe C, Barrett R, Cloetens P, Efficient concentration of high-energy x-rays for diffraction-limited imaging res-olution, Optica 4, 492–495, 2017.

14. Demidchik V, Shabala S, Coutts K, Tester M & Davies J. (2003). Free oxygen radicals regulate plasma membrane Ca2+- and K+-permeable channels in plant root cells. Journal of Cell Science, 1;116(1):81–8.

15. Desai M, Haigh M & Walkington H. (2018). Phytoremediation: Metal decontamination of soils after the sequential forestation of former opencast coal land. Science of the Total Environment, 15;656:670–680. doi: 10.1016/j.scitotenv.

16. Dierolf M, Menzel A, Thibault P, Schneider P, Kewish C, Wepf R, Bunk O & Pfeiffer F. (2010). Ptychographic X-ray computed tomography at the nanoscale. Nature, 467: 436–439.

17. Durán P, Viscardi S, Acuña J, Cornejo, Rosario Azcón P & de la Luz Mora M. (2018). Endophytic selenobacteria and arbuscular mycorrhizal fungus for Selenium biofortification and Gaeumannomyces graminis biocontrol. Journal of soil science and plant nutrition, 18(4) Temuco dic.

18. Esitken A. (1970). Use of Plant Growth Promoting Rhizobacteria in Horticultural Crops. In book: Bacteria in Agrobiology: Crop Ecosystems, Chapter 8. pages 189–235.

19. Furuta K, Yadav S, Lehesranta S, Belevich I, Miyashima S, Heo J, Vatén A, Lindgren O, De Rybel B, Van Isterdael G, Somervuo P, Lichtenberger R, Rocha R, Thitamadee S, Tähtiharju S, Auvinen P, Beeckman T, Jokitalo E & Helariutta Y. (2014). Arabidopsis NAC45/86 direct sieve element morphogenesis culminating in enucleation. Science, 345(6199): 933–937. DOI: 10.1126/science.1253736

20. Guizar-Sicairos M, Boon J, Mader K, Diaz A, Menzel A & Bunk O. (2015). Quantitative interior x-ray nanotomography by a hybrid imaging technique. Optica, 2(3): 259–266. https://doi.org/10.1364/OPTICA.2.000259

21. Guizar-Sicairos M, Diaz A, Holler M, Lucas M, Menzel A, Wepf R & Bunk O. (2011). Phase tomography from x-ray coherent diffractive imaging projections. Optics Express, 1(22): 21345–21357. https://doi.org/10.1364/OE.19.021345

22. Guizar-Sicairos M, Johnson I, Diaz A, Holler M, Karvinen P, Stadler H-C, Dinapoli R, Bunk O & Menzel A. (2014). High-throughput ptychography using Eiger: scanning X-ray nano-imaging of extended regions. Optics Express, 22(12):14859–14870. https://doi.org/10.1364/OE.22.014859

23. Gupta M & Gupta S. (2016). An overview of Selenium Uptake, Metabolism and Toxicity in plants, Frontiers in Plant Science, 2016(7):2074. doi: 10.3389/fpls.2016.02074.

24. Helin J, Ikonen A & Hjerpe T. (2010) Review of element specific data for biosphere assessment BSA-2009. Working Report 2010-37, Posiva Oy.

25. Holler M, Raabe J, Wepf R, Shahmoradian SH, Diaz A, Sarafimov B, Lachat T, Walther H & Vitins M (2017). OMNY PIN — A versatile sample holder for tomographic measurements at room and cryogenic temperatures. Review of Scientific Instruments, 88(11): doi:10.1063/1.4996092

26. Holler M, Raabe J, Diaz A, Guizar-Sicairos M, Wepf R, Odstrcil M, Shai F, Panneels V, Menzel A, Sarafimov B, Maag S, Wang X, Thominet V, Walther H, Lachat T, Vitins M & Bunk O. (2018). OMNY—A tOMography Nano crYo stage. Review of Scientific Instruments, 89, 043706. https://doi.org/10.1063/1.5020247

27. Hu P, Wang Y, Przybyłowicz W, Li Z, Barnabas A, Wu L, Luo Y & Mesjasz-Przybyłowicz J. (2015). Elemental distribution by cryo-micro-PIXE in the zinc and cadmium hyperaccumulator Sedum plumbizincicola grown naturally. Plant and Soil. 388(1–2) 267–282.

28. Huang X, Yan H, Harder R, Hwu Y, Robinson I & Chu Y. (2014). Optimization of overlap uniformness for ptychography. Optics Express, 22(10):12634–12644. https://doi.org/10.1364/OE.22.012634

29. Jiang Y, El Mehdawi A, Tripti, Lima L, Stonehouse G, Fakra S, Hu Y, Qi H & Pilon-Smits E. (2018). Characterization of Selenium Accumulation, Localization and Speciation in Buckwheat–Implications for Biofortification. Frontiers in Plant Science, 2018; 9: 1583. doi: 10.3389/fpls.2018.01583

30. Khan S, Zada S, Ahmad S, Lv J & Fu P. (2019). Concurrent biomineralization of silver ions into Ag0 and AgxO by Leptolyngbya strain JSC-1 and the establishment of its axenic culture. Chemosphere, 215, Pages 693–702.

31. Krajcarová L, Novotný K, Kummerová M, Dubová J, Gloser V & Kaiser J. (2017). Mapping of the spatial distribution of silver nanoparticles in root tissues of Vicia faba by laser-induced breakdown spectroscopy (LIBS) Talanta. 173:28–35. doi.org/10.1016/j.talanta.

32. Kurusu T, Kuchitsu K, Nakano M, Nakayama Y & Iida H. (2013). Plant mechanosensing and Ca2+ transport. Trends in Plant Science, 18, ISSUE 4, P227–233, April 01, 2013. DOI: https://doi.org/10.1016/j.tplants.2012.12.002

33. Li H, McGrath S & Zhao F (2008). Selenium uptake, translocation and speciation in wheat supplied with selenate or selenite. New Phytology, 178: 92–102, doi: 10.1111/j.1469-8137.2007.02343.x

34. Lincoln C, Britton J & Estelle M. (1990). Growth and Development of the axr1 Mutants of Arabidopsis. The Plant Cell, Vol. 2, 1071–1 080

35. Lintern M, Anand R, Ryan C & Paterson D. (2013). Natural gold particles in Eucalyptus leaves and their relevance to exploration for buried gold deposits. Nature Communications, 4: 2614

36. Liu S, Hu Q, Luo S, Li Q, Yang X, Wang X & Wang S. (2015) Expression of wild-type PtrIAA14.1, a poplar Aux/IAA gene causes morphological changes in Arabidopsis. Frontiers in Plant Science, 02 June 2015. doi.org/10.3389/fpls.2015.00388

37. Longchamp M, Angeli N & Castrec-Rouelle M. (2016). Effects on the accumulation of calcium, magnesium, iron, manganese, copper and zinc of adding the two inorganic forms of selenium to solution cultures of Zea mays. Plant Physiology and Biochemistry. 98:128–37. doi: 10.1016/j.plaphy.2015.11.013.

38. Lusa M, Bomberg M, Aromaa H, Knuutinen & Lehto J. (2015). The microbial impact on the sorption behaviour of selenite in an acidic, nutrient-poor boreal bog. Journal of Environmental Radioactivity, 147: 85–96.

39. Lusa M, Help H, Honkanen AP, Parkkonen J, Kalasova D & Blomberg M. (2019). Bacterial effects on selenium uptake in plants. Environmental Research. Available online 13 August 2019, 108642. https://doi.org/10.1016/j.envres.2019.108642

40. Lusa M, Knuutinen J & Blomberg M. (2017). Uptake and reduction of Se(IV) in two heterotrophic aerobic Pseudomonads strains isolated from boreal bog environment. AIMS Microbiology, 4: 798–814.

41. Lusa M, Lehto J, Aromaa H, Knuutinen J & Bomberg M. (2016) The Uptake of Radioiodide by Paenibacillus sp., Pseudomonas sp., Burkholderia sp. and Rhodococcus sp. isolated from a Boreal Nutrient-poor Bog. Journal of Environmental Sciences. 44, 26–37.

42. Mirone A, Brun E, Gouillart E, Tafforeau P & Kieffer J. (2014) The PyHST2 hybrid distributed code for high speed tomographic reconstruction with iterative reconstruction and a priori knowledge capabilities. Nuclear Instruments and Methods in Physics Research Section B: Beam Interactions with Materials and Atoms, 324: 41–48.

43. Monshausen G, Miller N, Murphy S & Gilroy S. (2011). Dynamics of auxin-dependent Ca2+ and pH signaling in root growth revealed by integrating high-resolution imaging with automated computer vision-based analysis. Plant Journal. 65(2):309–18. doi: 10.1111/j.1365-313X.2010.04423.x.

44. Morawe Ch, Barrett R, Cloetens P, Lantelme B; Peffen J-Ch, Vivo A. (2015) Graded multilayers for figured Kirkpatrick-Baez mirrors on the new ESRF end station ID16A. Proc. SPIE, 9588, 958803.

45. Odstrcil M. et al, manuscript in preparation

46. Ojuederie O & Babalola O. (2017). Microbial and Plant-Assisted Bioremediation of Heavy Metal Polluted Environments. International Journal of Environmental Research and Public Health, 4;14(12). pii: E1504. doi: 10.3390/ijerph14121504.e4

47. Ondrasek G, Rengel Z, Clode P, Kilburn M, Guagliardo P & Romic D. (2019). Zinc and cadmium mapping by NanoSIMS within the root apex after short-term exposure to metal contamination. Ecotoxicology and Environmental Safety. 171:571–578. doi.org/10.1016/j.ecoenv.

48. Paganin D, Mayo S C, Gureyev T E, Miller P R, Wilkins S W, Simultaneous phase and amplitude extraction from a single defocused image of a homogeneous object, J. Microsc. 206, 33 (2002)

49. Perrot-Rechenmann C. (2010). Cellular Responses to Auxin: Division versus Expansion. Cold Spring Harbor Perspectives in Biology, 2(5):a001446. doi: 10.1101/cshperspect.a001446

50. Pirozzi N, Hoogenboom J & Giepmans B (2018). ColorEM: analytical electron microscopy for element-guided identification and imaging of the building blocks of life. Histochemistry and Cell Biology. 150,(5): 509–520.

51. Reynolds R & Pilon-Smits E. (2018). Plant selenium hyperaccumulation-Ecological effects and potential implications for selenium cycling and community structure. Biochimica et Biophysica Acta (BBA) - General Subjects. 1862(11): 2372–2382.

52. Rodrigues E, Gomes M, Duran N, Cassanji J, da Cruz T, Sant’Anna Neto A, Savassa S, de Almeida E & Carvalho H. (2018). Laboratory Microprobe X-Ray Fluorescence in Plant Science: Emerging Applications and Case Studies. Frontiers in Plant Science, 9: 1588. doi: 10.3389/fpls.2018.01588

53. Shahmoradian S, Tsai E, Diaz A, Guizar-Sicairos M, Raabe J, Spycher L, Britschgi M, Ruf A, Stahlberg H & Holler M. (2017). Three-Dimensional Imaging of Biological Tissue by Cryo X-Ray Ptychography. Scientific Reports, 24; 7(1):6291.

54. Singh P, Mohanta T & Sinha A. (2015). Unraveling the Intricate Nexus of Molecular Mechanisms Governing Rice Root Development: OsMPK3/6 and Auxin-Cytokinin Interplay. PLOS ONE, DOI:10.1371/journal.pone.0123620

55. Solé V, Papillon E, Cotte M, Walter Ph & Susini J. (2007). A multiplatform code for the analysis of energy-dispersive X-ray fluorescence spectra. Spectrochimica Acta Part B: Atomic Spectroscopy, 62(1): 63–68.

56. Staicu L, van Hullebusch E & Lens P. (2017). Wastewaters and Physical–Chemical Treatment Technologies. Bioremediation of Selenium Contaminated Wastewater pp 103–130 | Industrial Selenium Pollution:

57. Tangahu B, Abdullah S, Basri H, Idris M, Anuar N & Mukhlisin M. (2011). A Review on Heavy Metals (As, Pb, and Hg) Uptake by Plants through Phytoremediation. International Journal of Chemical Engineering. 2011; 939161. doi.org/10.1155/2011/939161

58. Thibault P, Dierolf M, Menzel A, Bunk O, David C & Pfeiffer F. (2008). High-Resolution Scanning X-ray Diffraction Microscopy. Science, 321(5887): 379–382. DOI: 10.1126/science.1158573

59. Thibault P & Guizar-Sicairos M. (2012). Maximum-likelihood refinement for coherent diffractive imaging. New Journal of Physics, Volume 14.

60. Tran T, Zhou F, Yang W, Wang M, Dinha Q, Wang D & Liang D. (2018). Detoxification of mercury in soil by selenite and related mechanisms. Ecotoxicology and Environmental Safety, 159, Pages 77–84

61. van Heel M & Schatz M. (2005). Fourier shell correlation threshold criteria. Journal of Structural Biology, 151 (3): 250–262. doi.org/10.1016/j.jsb.2005.05.009

62. Wang P, Menzies N, Lombi E, McKenna B, de Jonge M, Donner E, Blamey F, Ryan C, Paterson D, Howard D, James S & Kopittke P. (2013). Quantitative determination of metal and metalloid spatial distribution in hydrated and fresh roots of cowpea using synchrotron-based X-ray fluorescence microscopy. Science of The Total Environment. 463–464: 131–139.

63. Weinhardt V, Chen JH, Ekman A, McDermott G, Le Gros MA and Larabell C. (2019). Imaging cell morphology and physiology using X-rays, Biochemical Society Transactions. BST20180036; DOI: 10.1042/BST20180036

64. Xi W, Gong X, Yang Q, Yu H & Liou YC. (2016). Pin1At regulates PIN1 polar localization and root gravitropism. Nature Communications. 2016/01/21/online, DOI: 10.1038/ncomms10430

65. Ximénez-Embún P, Alonso I, Madrid-Albarrán Y, Cámara C. (2004). Establishment of Selenium Uptake and Species Distribution in Lupine, Indian Mustard, and Sunflower Plants. Journal of Agricultural and Food Chemistry, 52: 832–838.

66. Yu Y, Fu P, Huang Q, Zhang J & Li H. (2019). Accumulation, subcellular distribution, and oxidative stress of cadmium in Brassica chinensis supplied with selenite and selenate at different growth stages. Chemosphere, 216: 331–340

67. Zhao FJ, Moore K, Lombi E, Zhu YG. (2014). Imaging element distribution and speciation in plant cells. Trends in Plant Science. 19(3): 183–192

68. Zhao Y, Christensen S, Fankhauser C, Cashman J, Cohen J, Weigel D & Chory J. (2001). A Role for Flavin Monooxygenase–Like Enzymes in Auxin Biosynthesis. Science, 291; 306–309.

