## Extended Figures for manuscript for "High-resolution synchrotron imaging studies of intact fresh roots reveal soil bacteria promoted bioremediation and bio-fortification"

### EXTENDED MATERIAL LEGENDS

**Extended Figure 1: Electron densities of cryo- PXCT imaged roots**

These 3 graphs represent the electron density measurements from control, Se supplemented and Se+bacteria supplemented roots imaged using cryo-PXCT. For each average value shown in the attached graphs, >30 points across the samples were measured and averaged. Minimum (light blue bars), maximum (orange bars), median (light yellow bars) and mean (dark yellow bars) represent the electron densities of cell walls, cytoplasm and organelles, vacuole fluid and membranes. Error bars are included.

**Extended Figure 2: XRNF data for Se. Fe. Cr. Br and Ca**

While most chemical elements showed little difference in signal levels between different treatments, Se, Fe, Cr and Br showed different accumulation levels in roots, of which Se and Fe were most noticeable. Ca is presented as an element that did not change between treatments. Zn measurements with clear aggregates are shown in the bottom panel.

**Extended Figure 3: Example of Se measurements at organelle level in Se+bacteria supplanted root**

Se levels measured at organelle level, here shown in Se+bacteria supplemented root #1, of which individual cell types and organelles were most distinguishable. Small yellow boxes in the top panels represent regions from which Se was measured in cortex cells. Left, cytoplasm and ER, brightest signal corresponds to highest Se levels, darkest signal to low Se levels. Middle panel shows vacuole membranes from which signals were measured, and right panel cell vacuolic fluid areas. Highest Se signal levels were measured in cytoplasm (and ER) in all analyzed cell types; epidermis (brown color in insert picture), cortex (yellow) and endodermis (pink).The bars on the graph do not show error bars as they represent averaged values from multiple measurement points (Extended Figure 8). In all measured tissues, Se levels are highest cytoplasm (including ER).

**Extended Figure 4: Damaged root epidermal cells show increased Ca signal**

The XNH 3D reconstruction of a Se+bacteria supplemented root sample #2 on the left shows rod like Pseudomonas bacteria visible on the root surface. The cells marked with red arrow are also visibly damaged, and correspondingly show higher Ca signal in fluorescent map (top panel on the right). The Ca signal on damaged areas was ~2,5-fold stronger than in intact cell walls, and is shown in table. 17 individual points/areas were measured for both damaged and intact root epidermis values (n=17 + 17).

**Extended Figure 5: Root density plots for Se and Fe visualization of plotted areas**

Voxel-wise concentrations of Fe vs. Se in XRNF reconstructions of the measured samples. Scatter plots A-E correspond to the control, Se 1, Se 2, Se+bac 1, and Se+bac2, respectively. Panel F presents the same data in a single figure together with 5 regions of interest (labelled 1-5). The XRNF voxels falling inside the marked rois are presented for each sample over the XNH data in the bottom part of the figure. Notably the outlier region 5 corresponds to the damaged cells of epidermis in Se+bacteria supplemented samples.

**Extended Figure 6: A single slice electron density image of a Se+bacteria treated root imaged using 3D cryo-PXCT**

Pseudo-color image of a Se+bacteria supplemented root imaged using PXCT. Vacuolic cell organelles can be distinguished by their yellow color, and their density is smaller than cytoplasm (orange), endoplasmatic reticulum (red), cell walls (red, pink and white) and vacuolic membranes (red membranes inside yellow areas). As was shown with fluorescence measurements Fe localizes primarily to root cell walls, especially in cell corners. These high density areas show white signal. Fluorescence data also showed that Se accumulates primarily inside the cells; in cytoplasm and ER (orange and red areas) and in vacuole membranes shown here in orange and red. The white aggregates marked with black hollow arrows are likely Zn or Ca aggregates. Orange organelles in the cytoplasm (marked with narrow arrows) are mitochondria and/or other small cell organelles. Most cells in different tissues exhibit similar anatomy and density features. What is noticeable, is that the resolution of cryogenic PXCT imaging at 80nm resolution allows detection and distinction of individual cellular organelles: mitochondria, plastids, nuclei, cytoplasmic fluid, ER, vacuole membranes. The cells at the center of the root are vascular cells, and the ones with thicker cell walls (marked with asterisks) are phloem sieve elements.

**Extended Figure 7: Different root cell identities can be distiguished from PXCT reconstructions**

Panels on the left show electron density maps of a PXCT imaged control root, cross section (top panel) and longitudinal section (bottom panel). Panels on the right side are artificially colored to illustrate different cell layers normally present in roots (cross section at the top, longitudinal section in the middle, color legends at the bottom). Outermost cell layer is epidermis (brown), below it is cortex (yellow), endodermis (rosa), pericycle (purple), proto xylem (blue), metaxylem (light blue), procambium (light green), phloem companion cells (pink) and phloem sieve elements (red). All scale bars are 10 μm.

**Extended Figure 8: Overall root architectures of different treatments, growth arrested roots and abnormal lateral root anatomy of bacteria supplemented samples.**

This figure show overall architecture of imaged roots. Roots grown on normal media for 2 weeks (A), look similar to roots grown on normal media for 1 w and supplemented with Se for 1 weeks (B) whereas roots grown with Se+bacteria for 1 w (C) grow differently than agar controls; bacteria supplemented roots show defects in primary root elongation and branching. D is a close up of a control lateral root, E is a Se supplemented root and F , G and H show Se+bacteria supplemented lateral roots of similar age as D and E. Root in panel F exhibits stunted growth and swelling, and the roots in panel G and H show arrest in growth (dying meristem is white and opaque) and an extraordinary branching phenotype. In control samples, roots never emerge side by side. Scale bar in H is 100μm.

**Table 2: Electron densities of cell compartments**

Presented values of subcellular organelle density minimum, maximum, median and mean from different treatments are averages, each measured from n>30 areas around the sample. The size of the measured areas depended on the cell organelle size.

**Extended Figure 1: Electron densities of cryo- PXCT imaged roots**

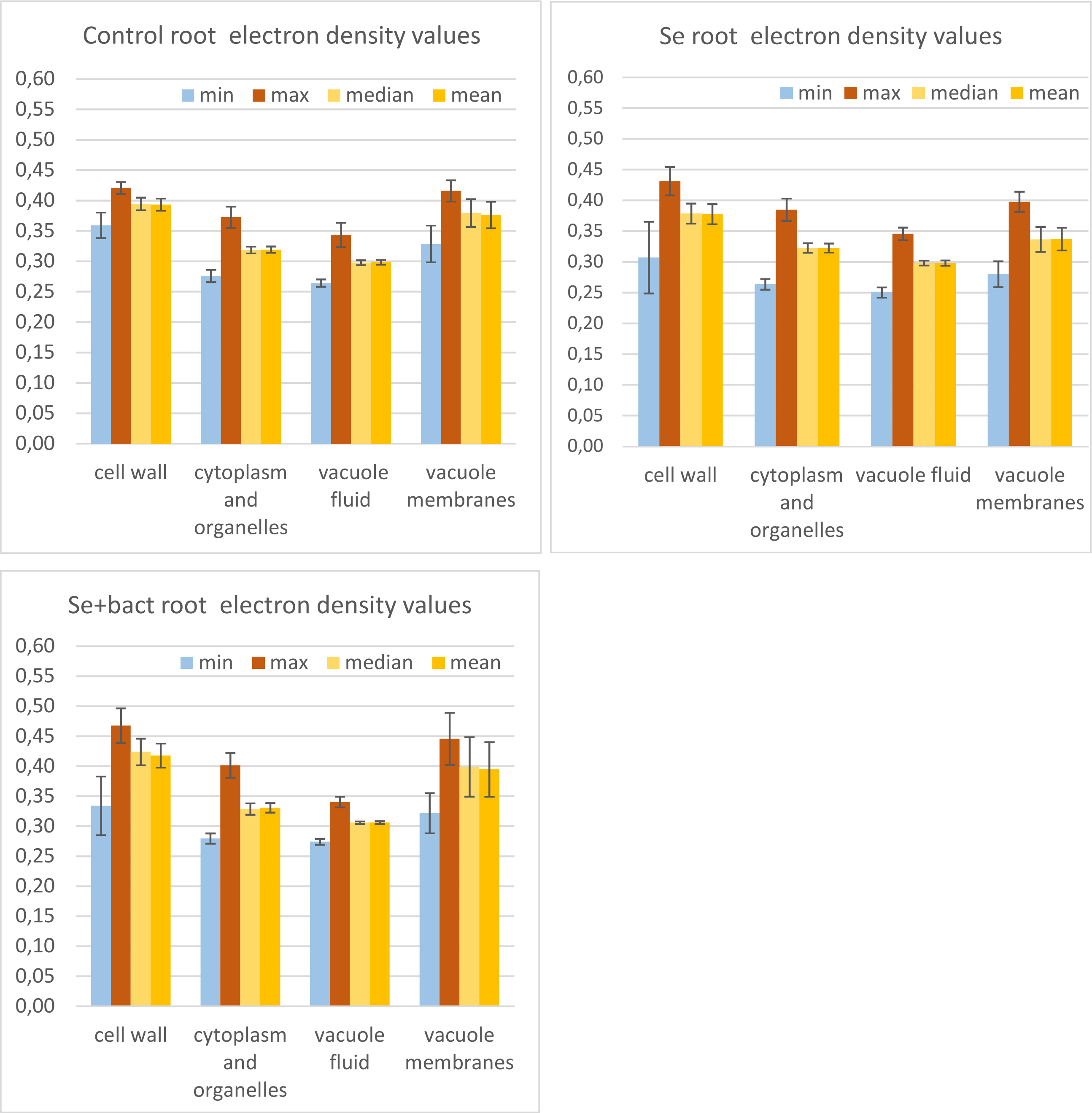

**Extended Figure 2: XRNF data for Se. Fe. Cr. Br and Ca**

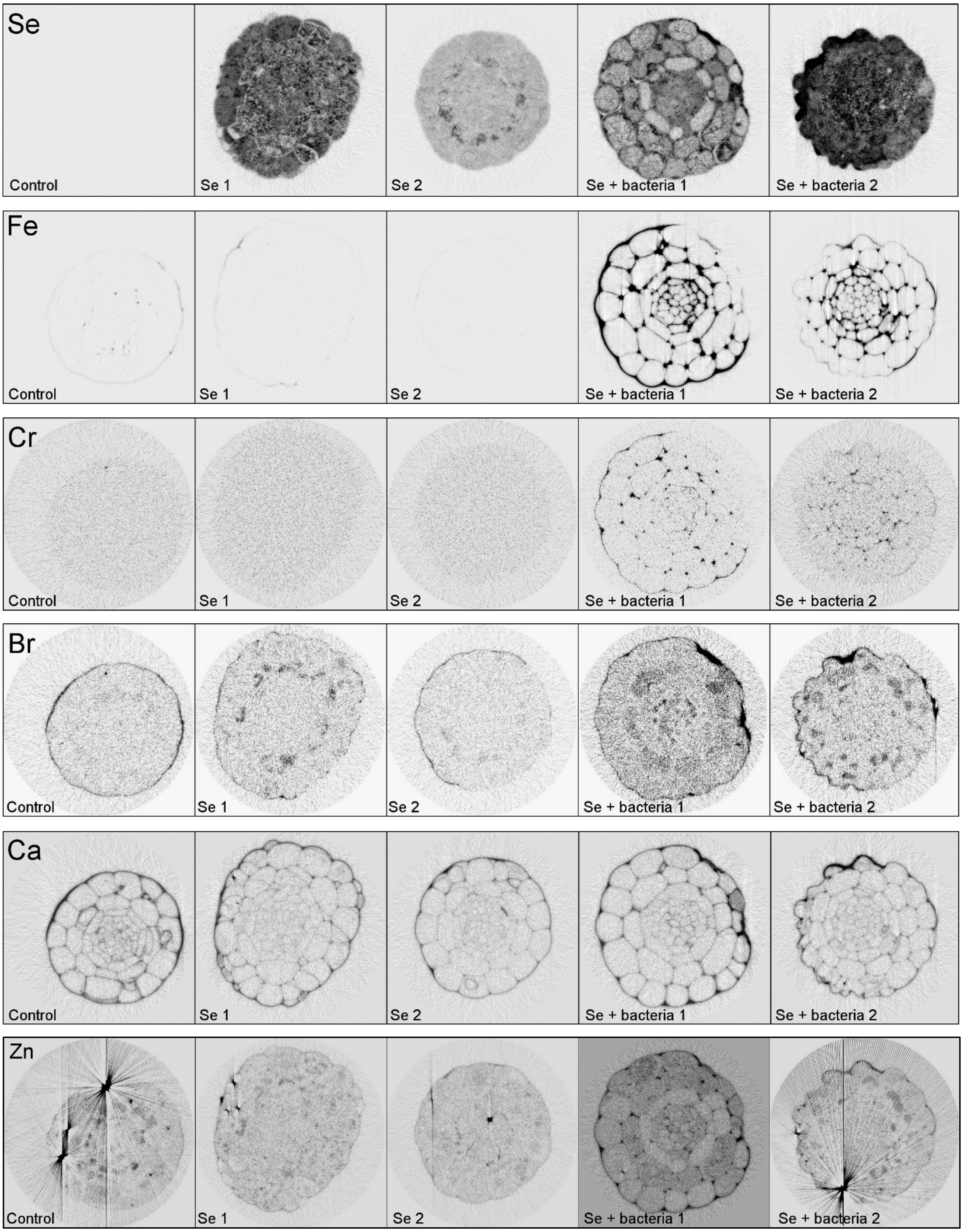

**Extended Figure 3: Example of Se measurements at organelle level in Se+bacteria supplemented root**

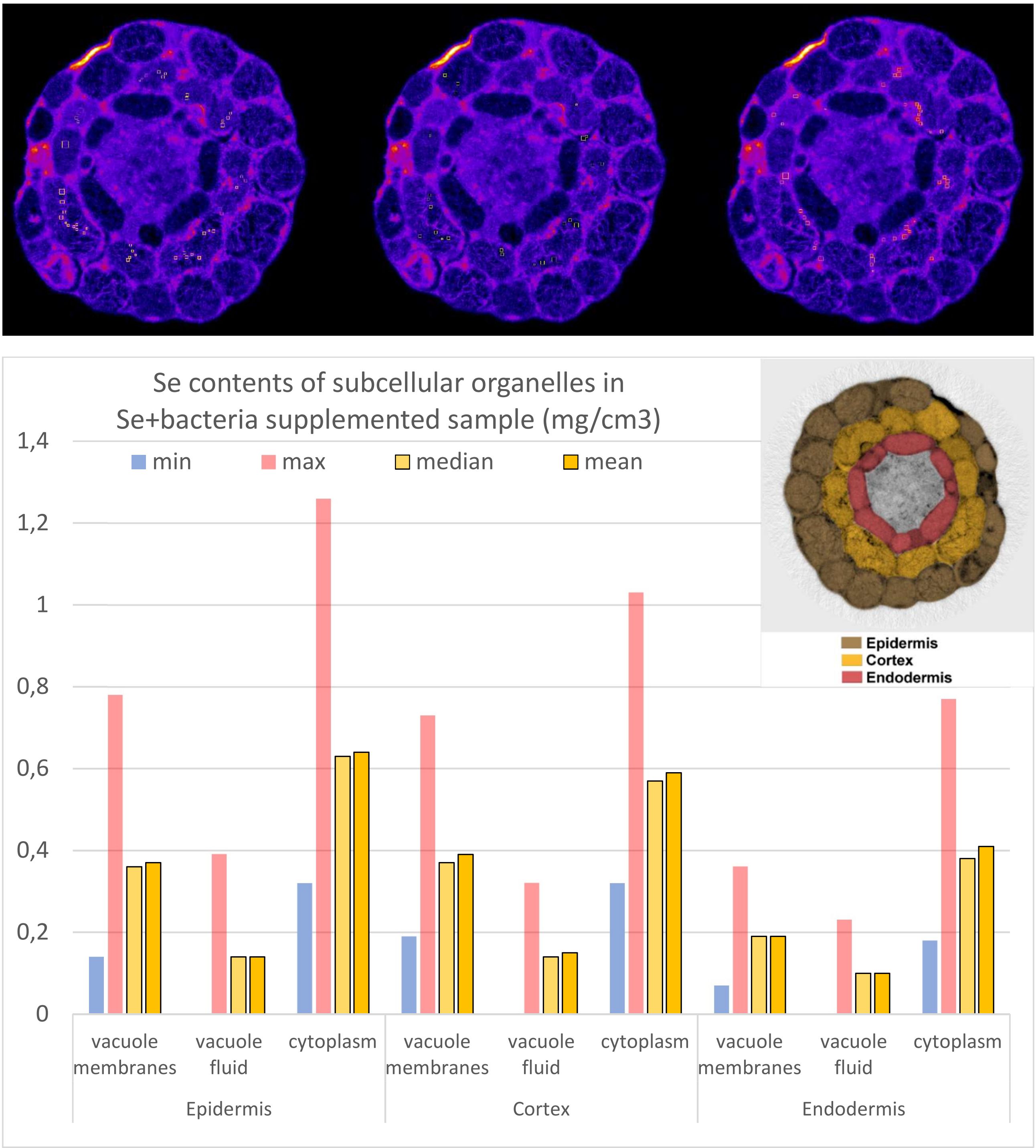

**Extended Figure 4: Damaged root epidermal cells show increased Ca signal**

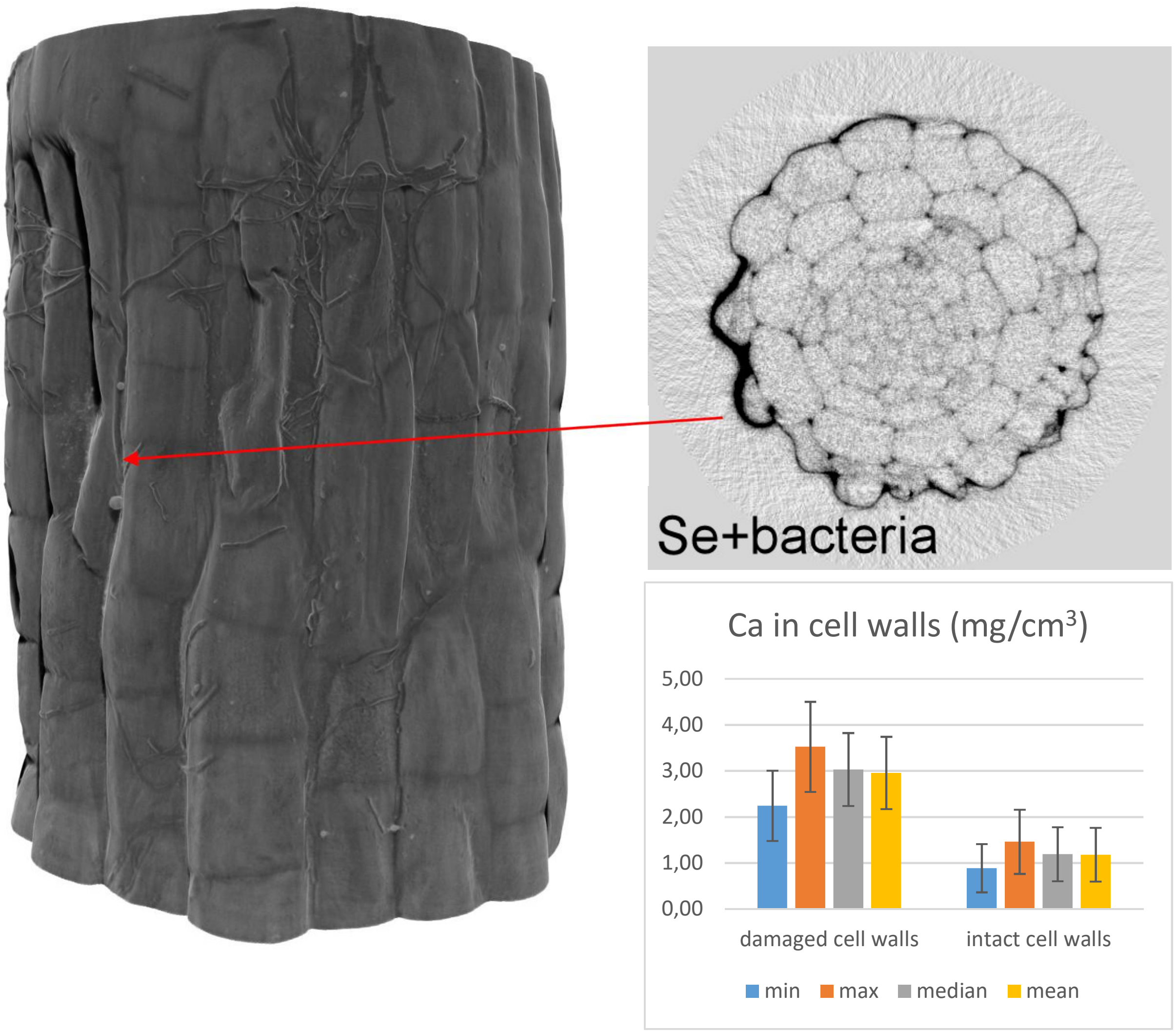

### Extended Figure 5 – Root density plots for Se and Fe visualization of plotted areas

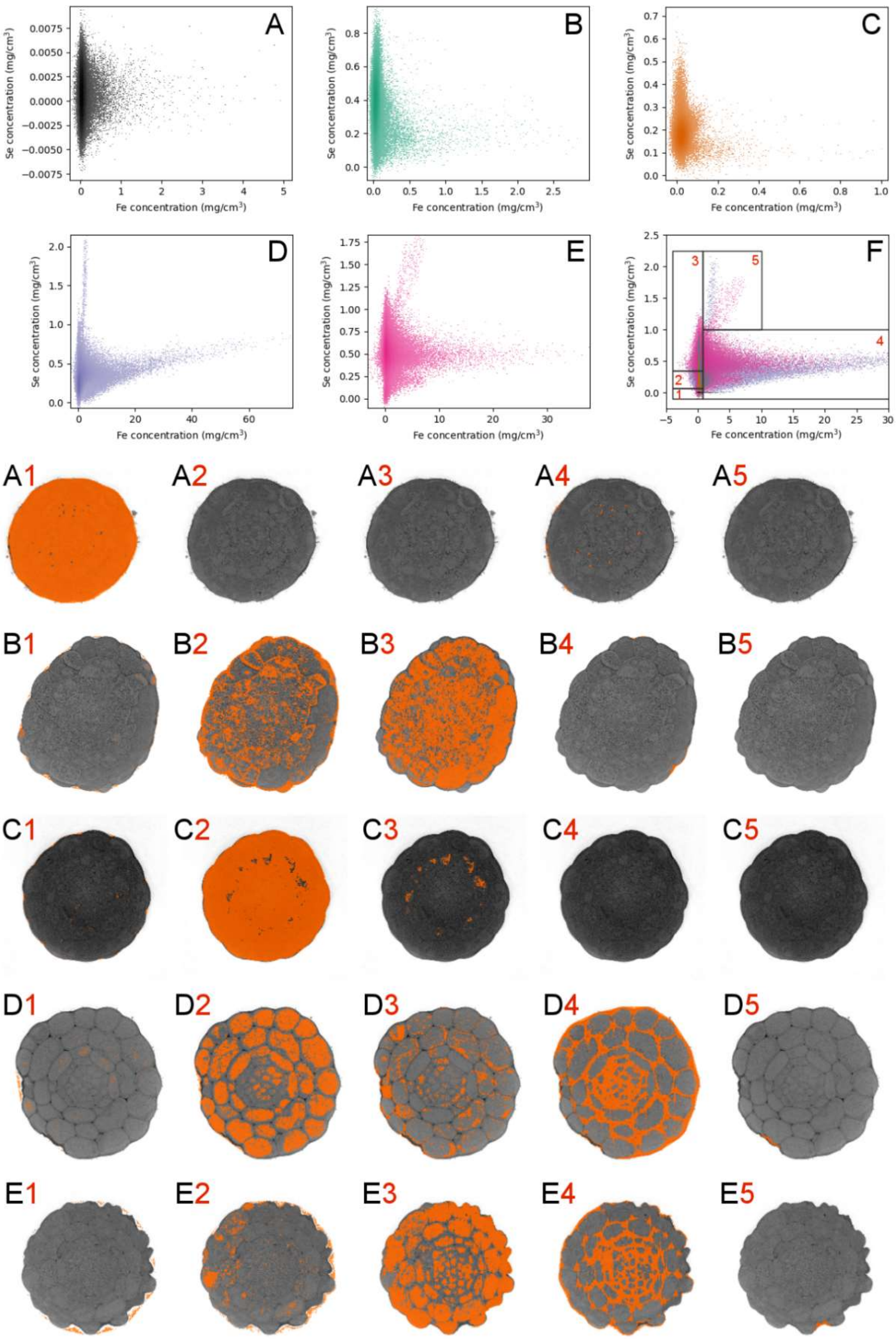

**Extended Figure 6: A single slice electron density image of a Se+bacteria treated root imaged using 3D cryo-PXCT**

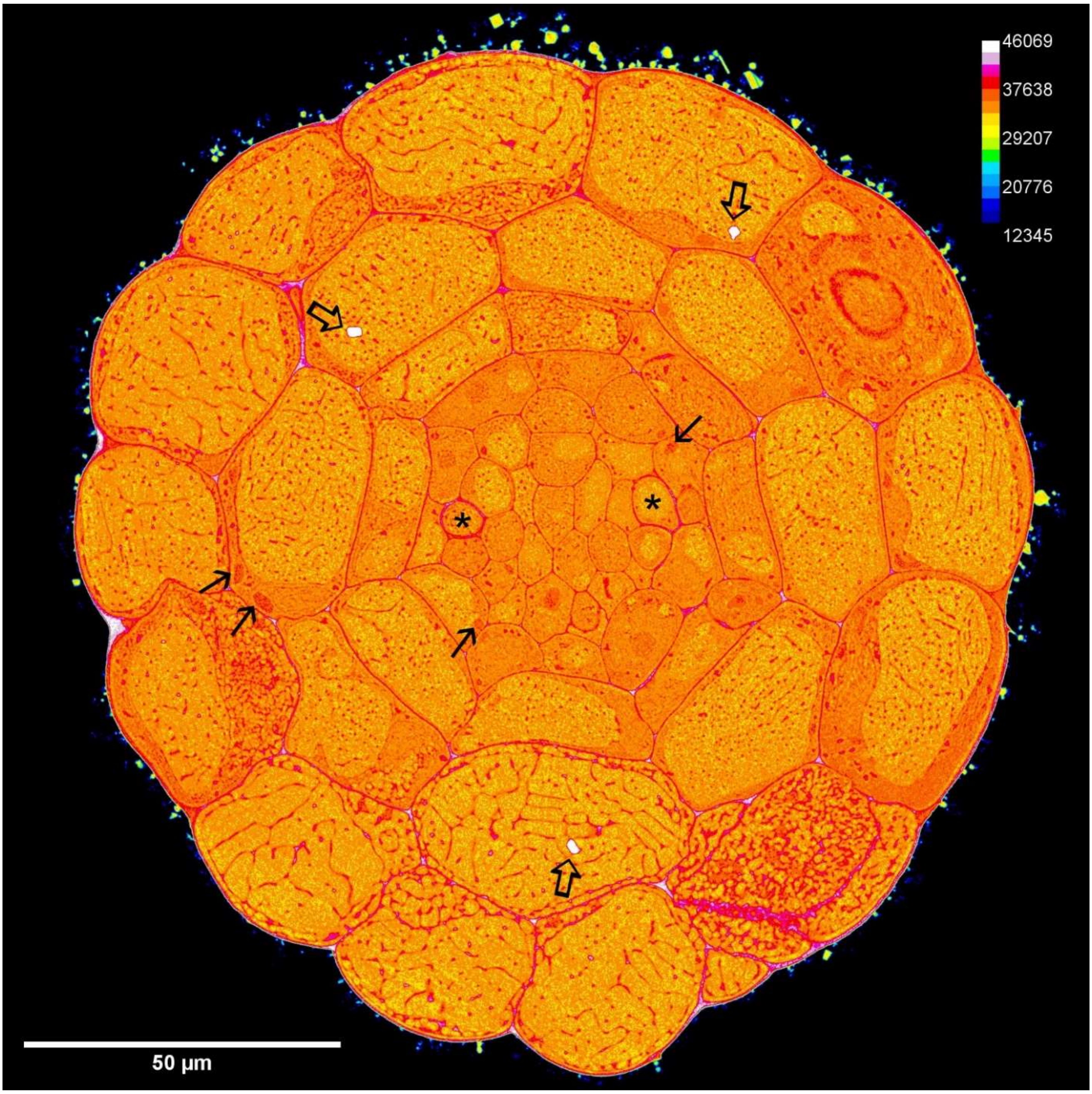

**Extended Figure 7: Different root cell identities can be distiguished from PXCT reconstructions**

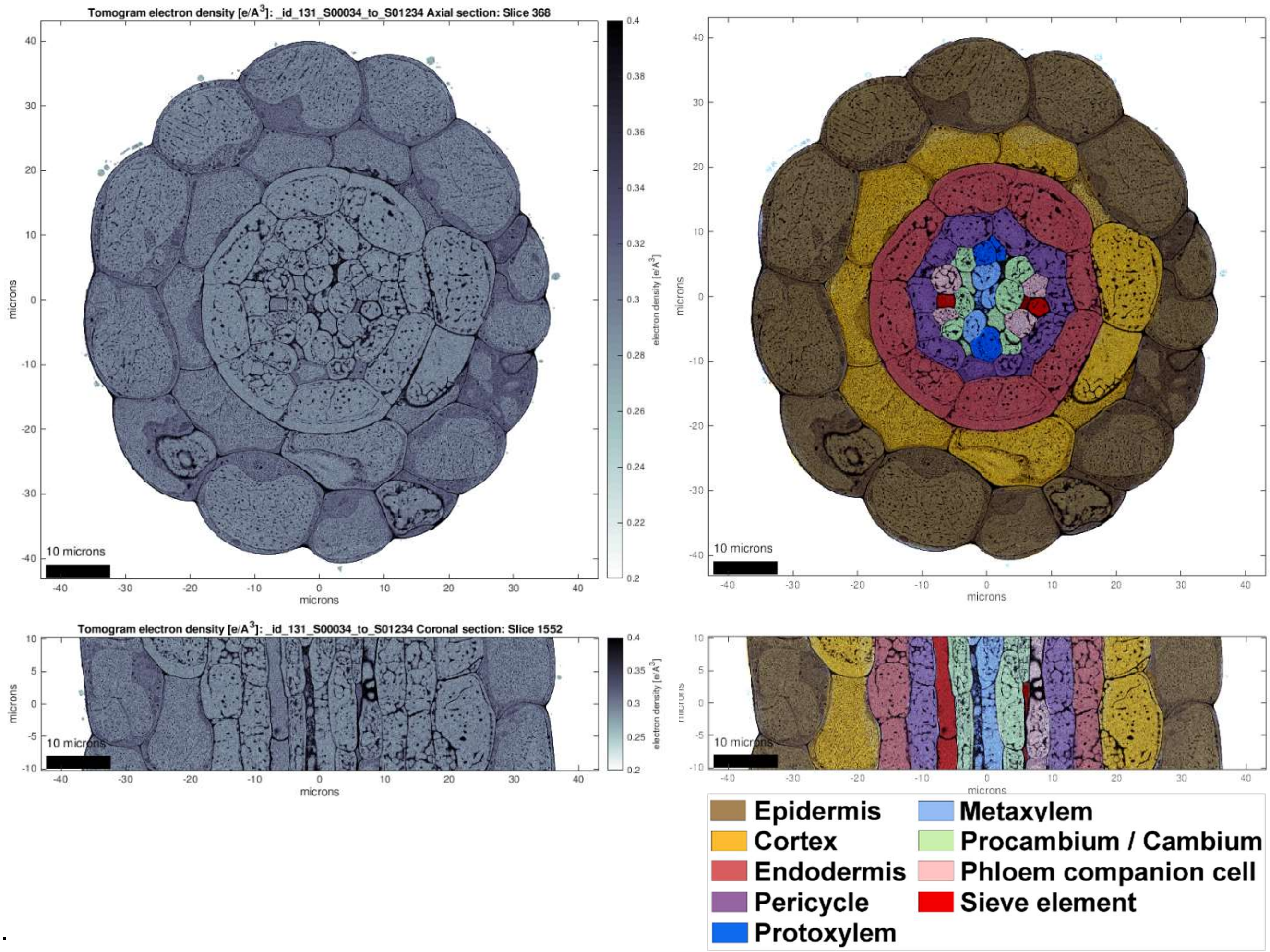


.

**Extended Figure 8: Overall root architectures of different treatments. growth arrested roots and abnormal lateral root anatomy of bacteria supplemented samples.**

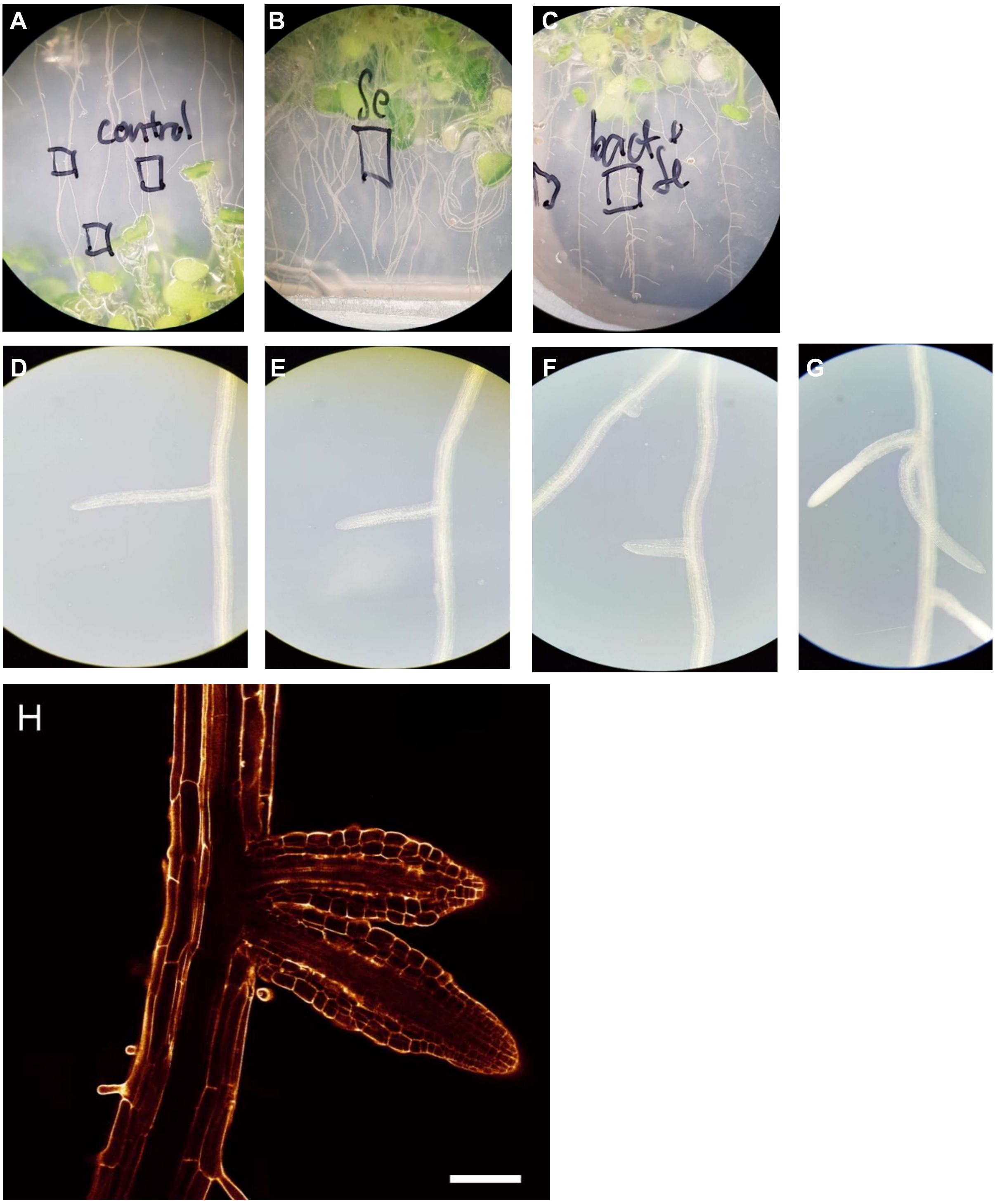

**Table 2: Overview of the cryo-PXCT measurement parameters**

| **Sample ID** | **id131** | **id132** | **id133** | **id134** | **id135** | **id136** |
|---|---|---|---|---|---|---|
| Name | control_s1 | bakt_sel_1 | sel_7 | bakt_sel_2 | control_2 | sel_9 |
| Field of view [µm$^2$] | 100 x 20 | 85 x 22 | 85 x 22 | 110 x 12 | 70 x 24 | 130 x 20 |
| Voxel size (nm) | 27.8 | 34.7 | 34.7 | 34.7 | 34.7 | 34.7 |
| # Projections | 1200 | 1200 | 1200 | 600 | 600 | 950 (1000) |
| Dose [MGy] | 2.5 | 15 | 15 | 6.5 | 7 | |
| Measurement time (projection) | 60 s | 54 s | 54 s | 41 s | 50 s | 72 s |
| Measurement time (total) | 22 h | 21 h | 21 h | 10 h | 10 h | |
| Sample diameter/ number of projections | 70 nm | 60 nm | 70 nm | 160 nm | 90 nm | 120 nm |
| FSC 3D | 90 nm | 80 nm | 90 nm | NA* | 124 nm | 130 nm |
| Figures | 2A, 2B, S1, S10 | 2C, S4 | | 2D | | |

Overview of the cryo-PXCT measurement parameters. (*) The FSC estimation for id134 is unrealistically low due to the insufficient amount of projections in each half-tomogram, given the large diameter of this sample.
