## Supplementary figures and images for "High-resolution synchrotron imaging studies of intact fresh roots reveal soil bacteria promoted bioremediation and bio-fortification"

### control root image 1

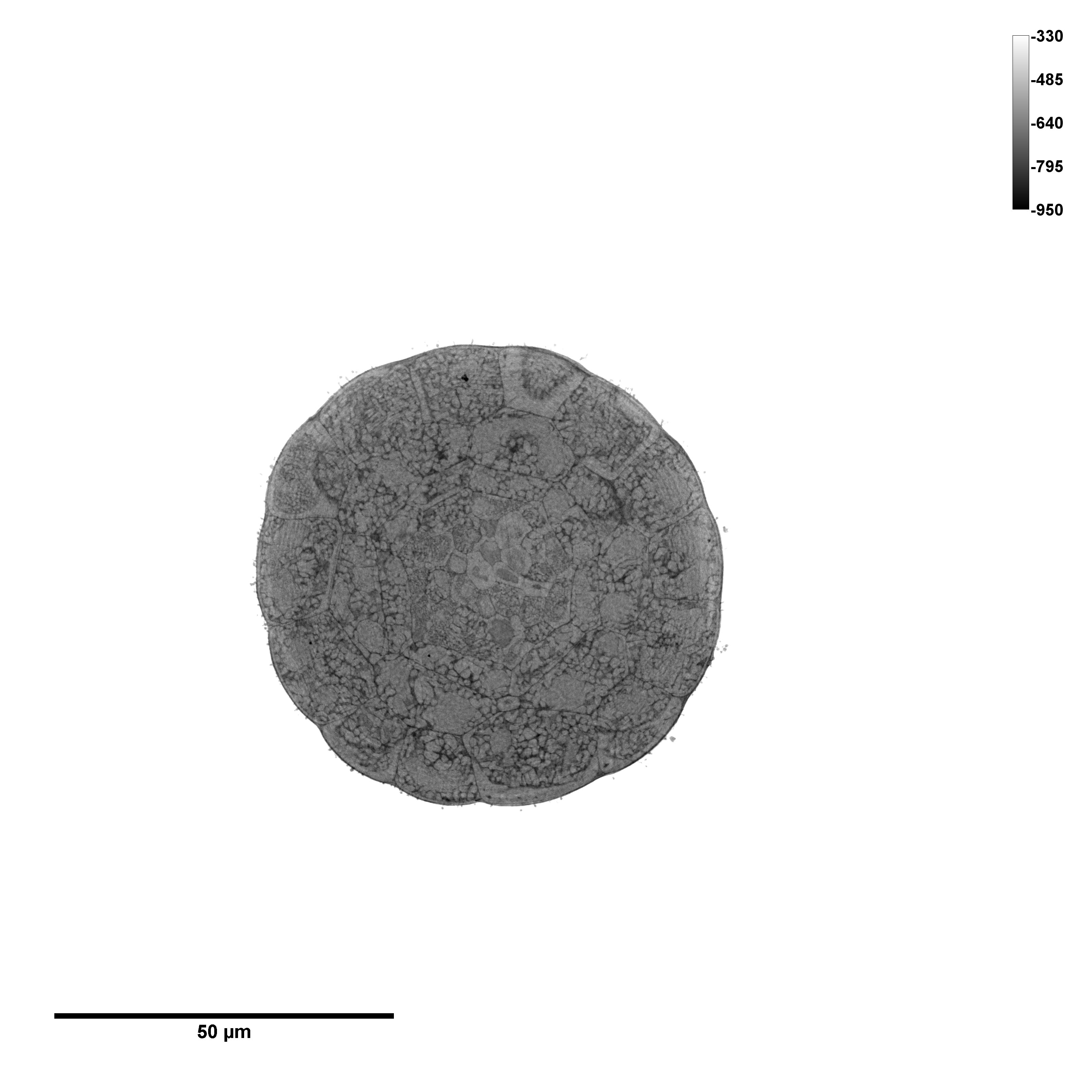

### control root image 2

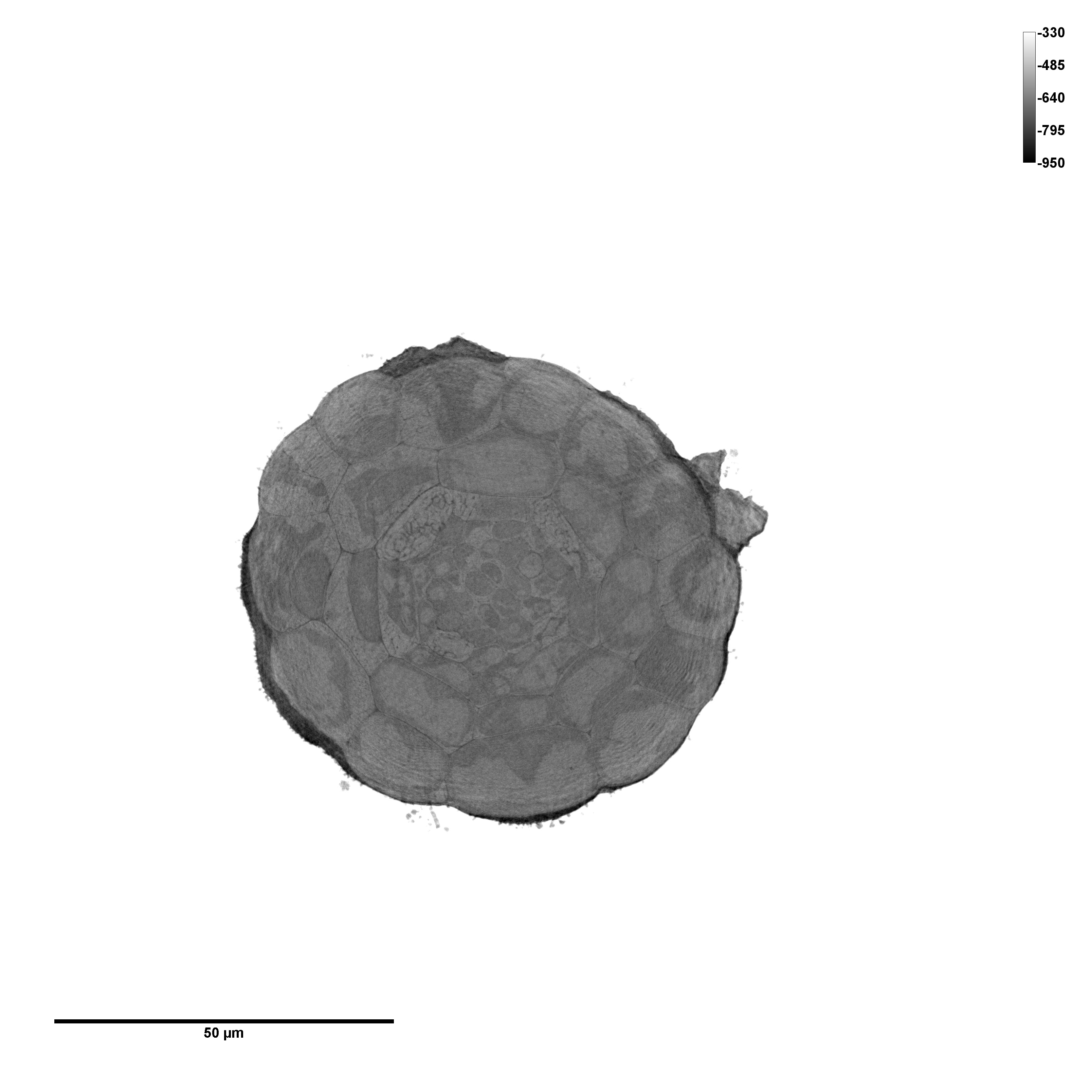

### Se + bact root 1 image 1

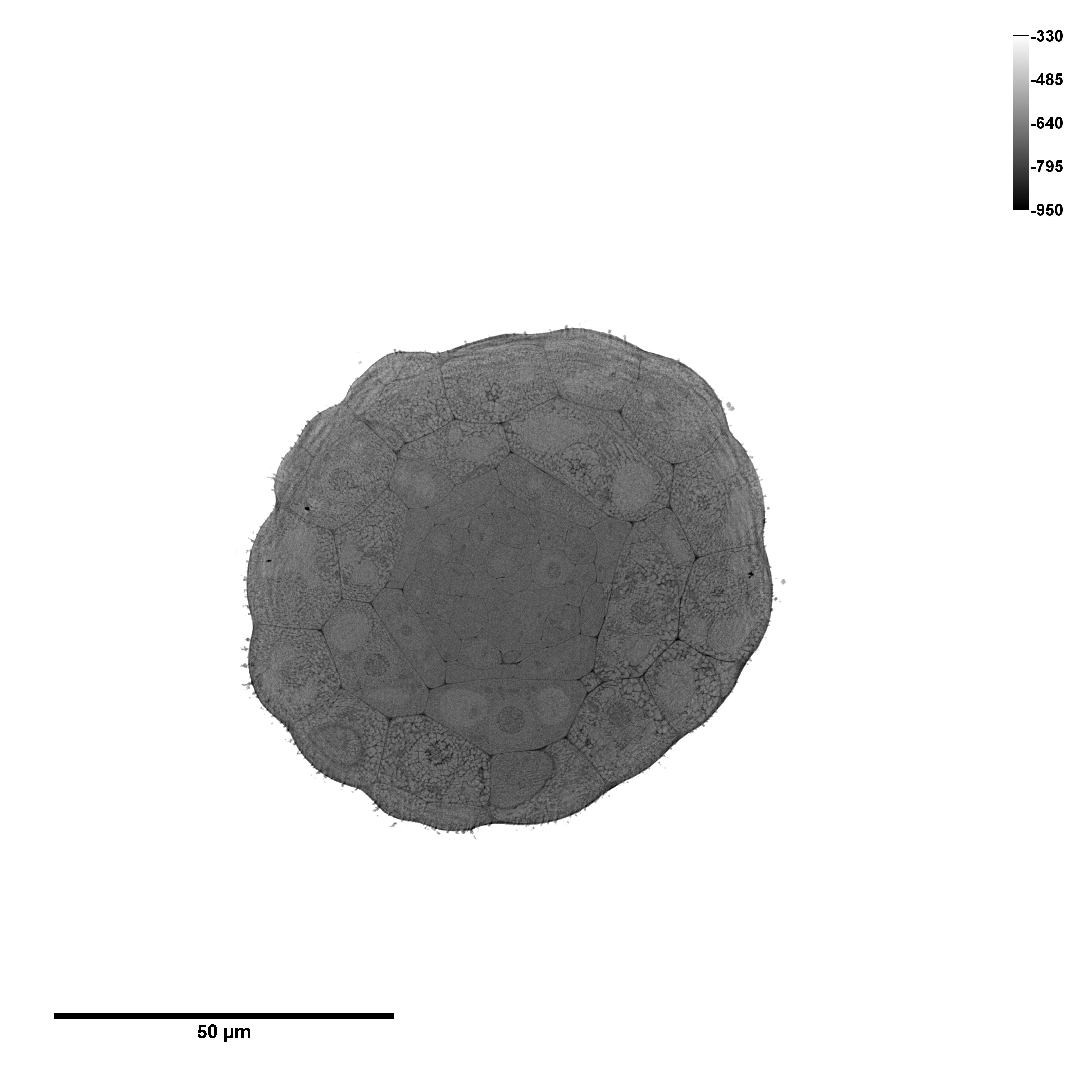

### Se + bact root 1 image 2

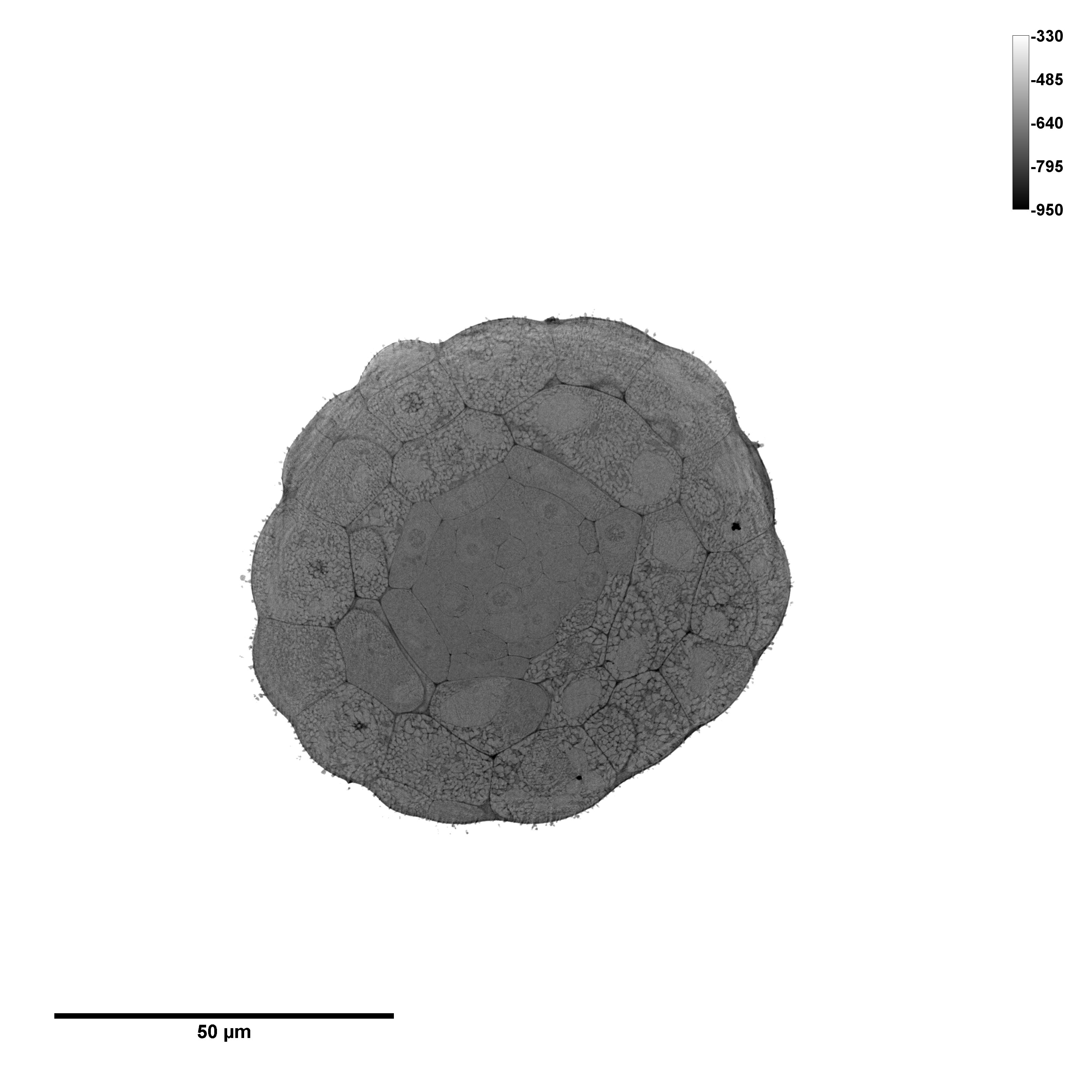

### Se + bact root 2 image 1

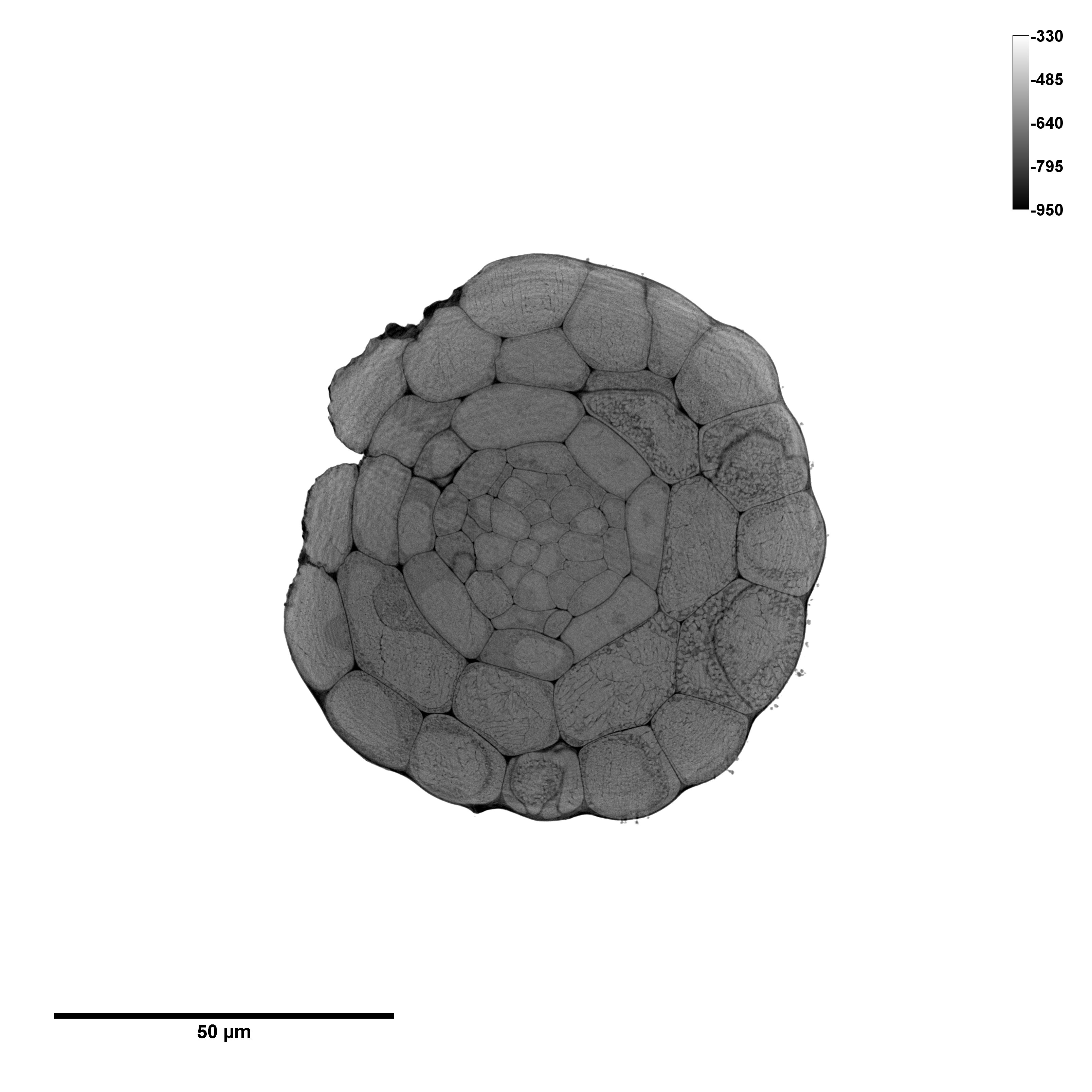

### Se + bact root 2 image 2

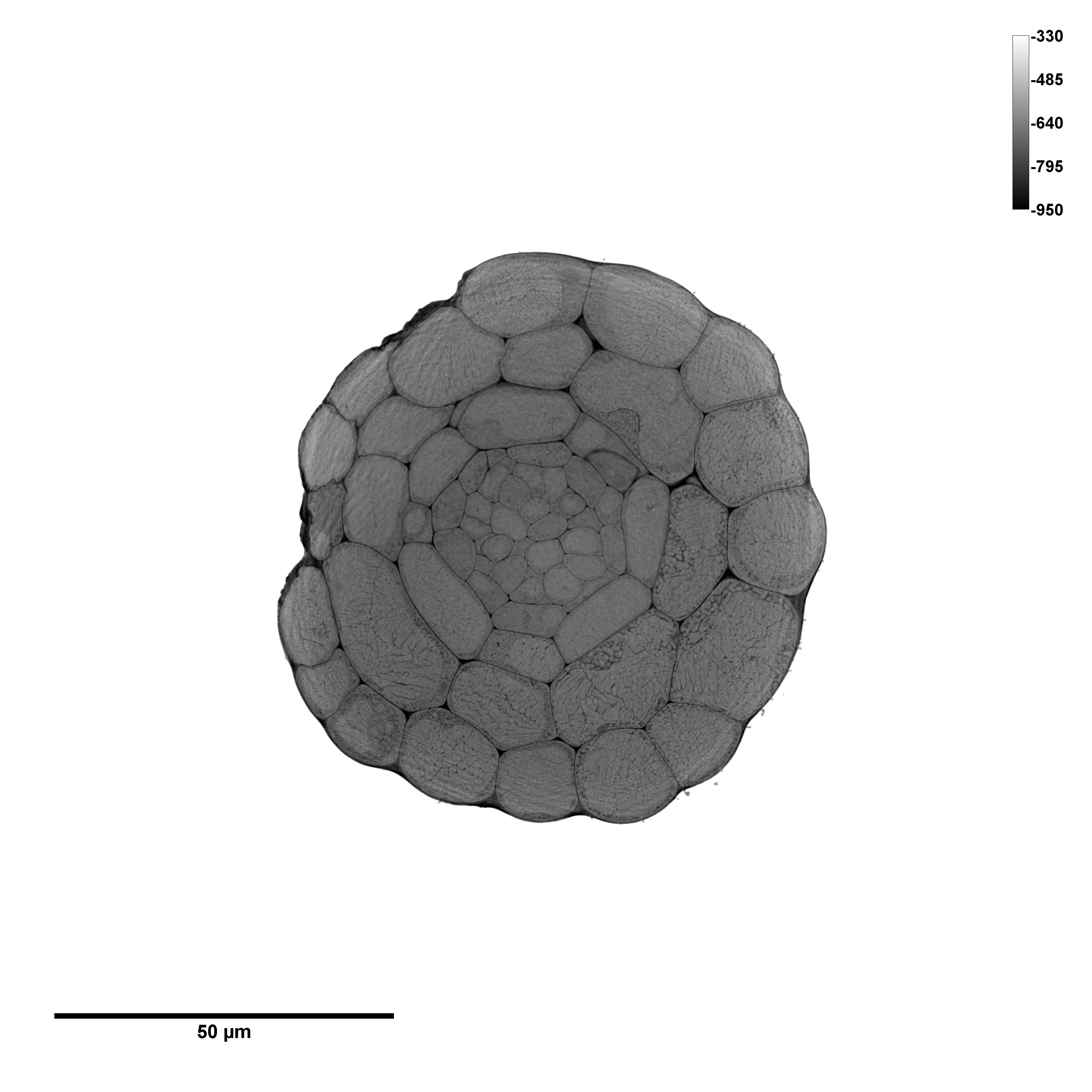

### Se + bact root 2 image 3

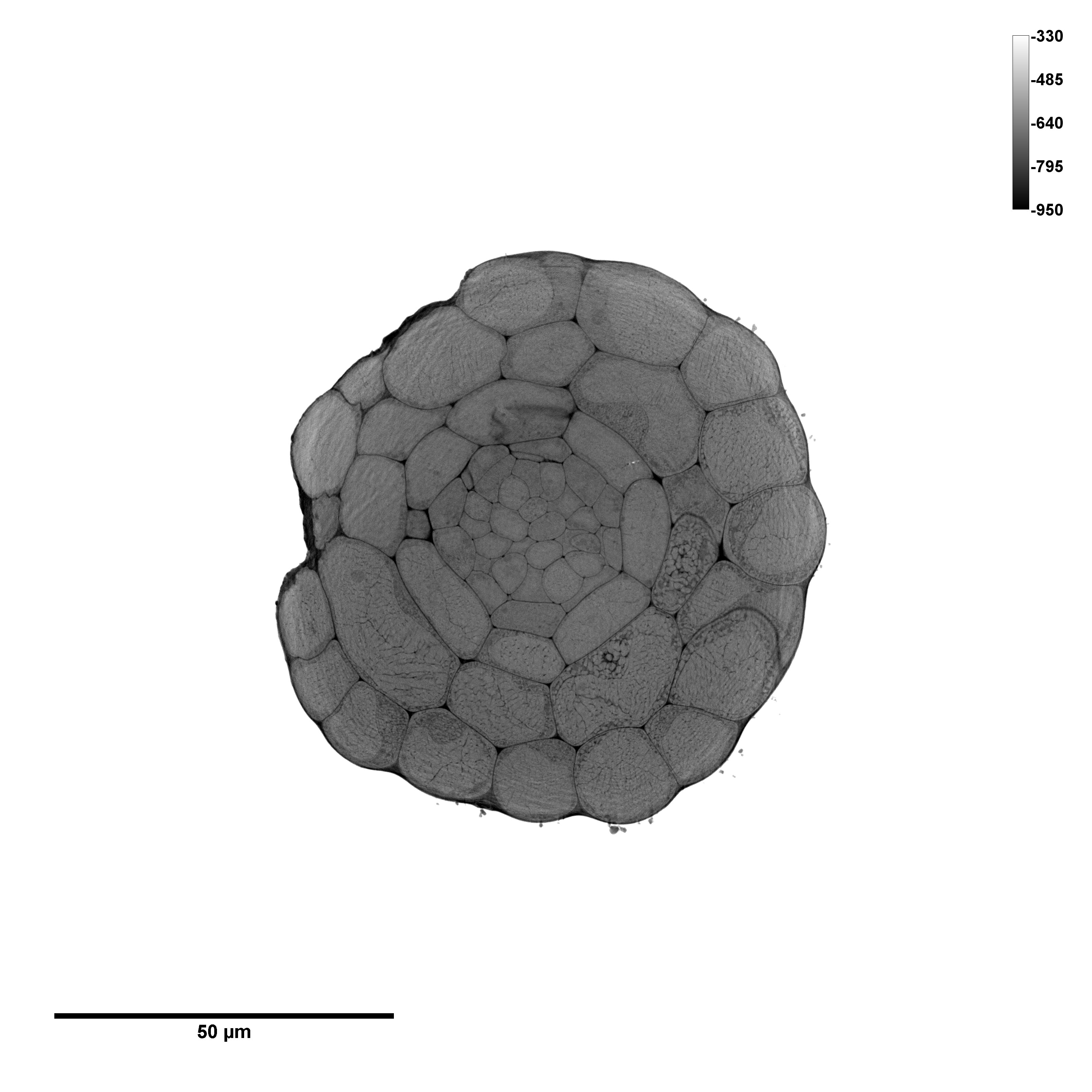

### Se + bact root 3 image 1

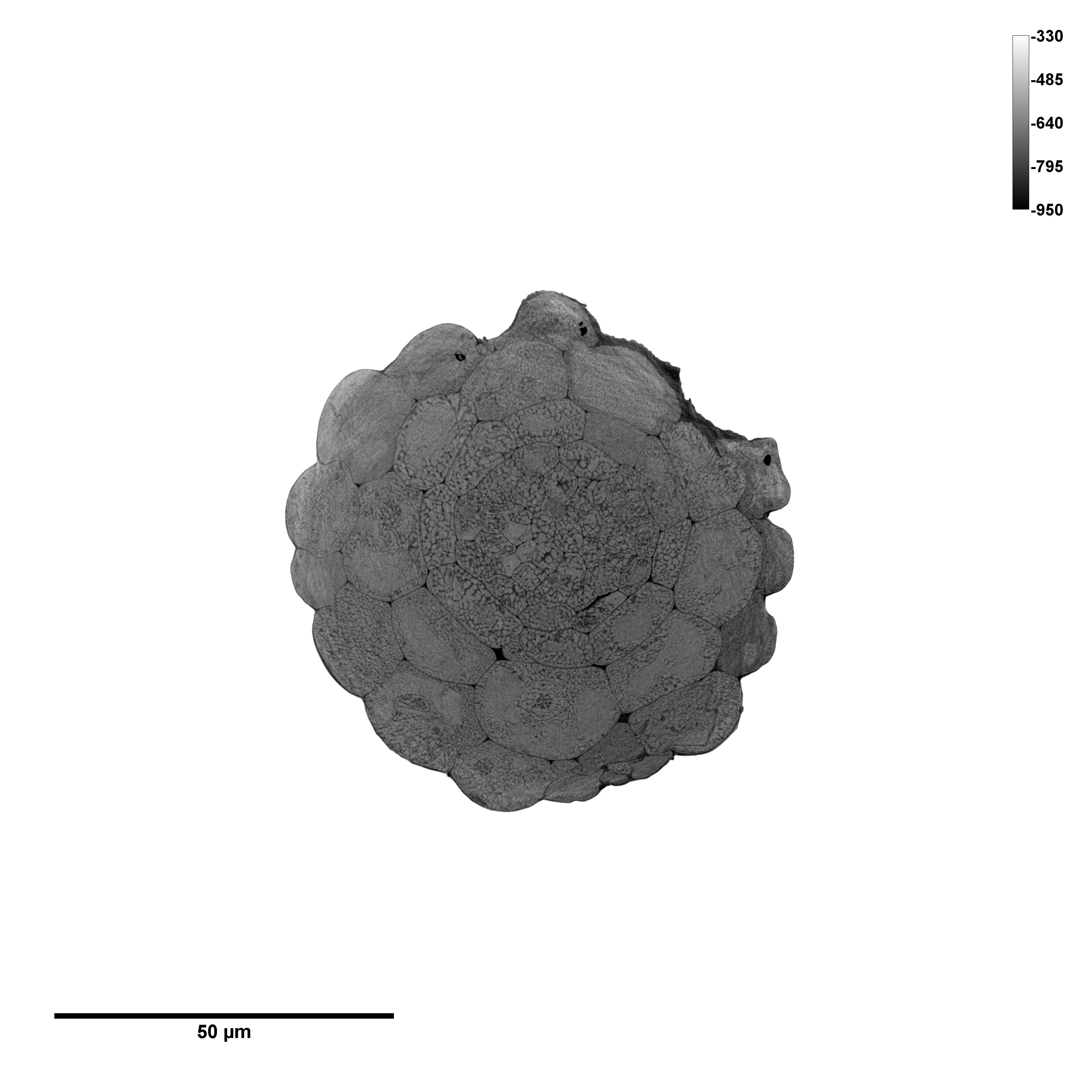

### Se + bact root 3 image 2

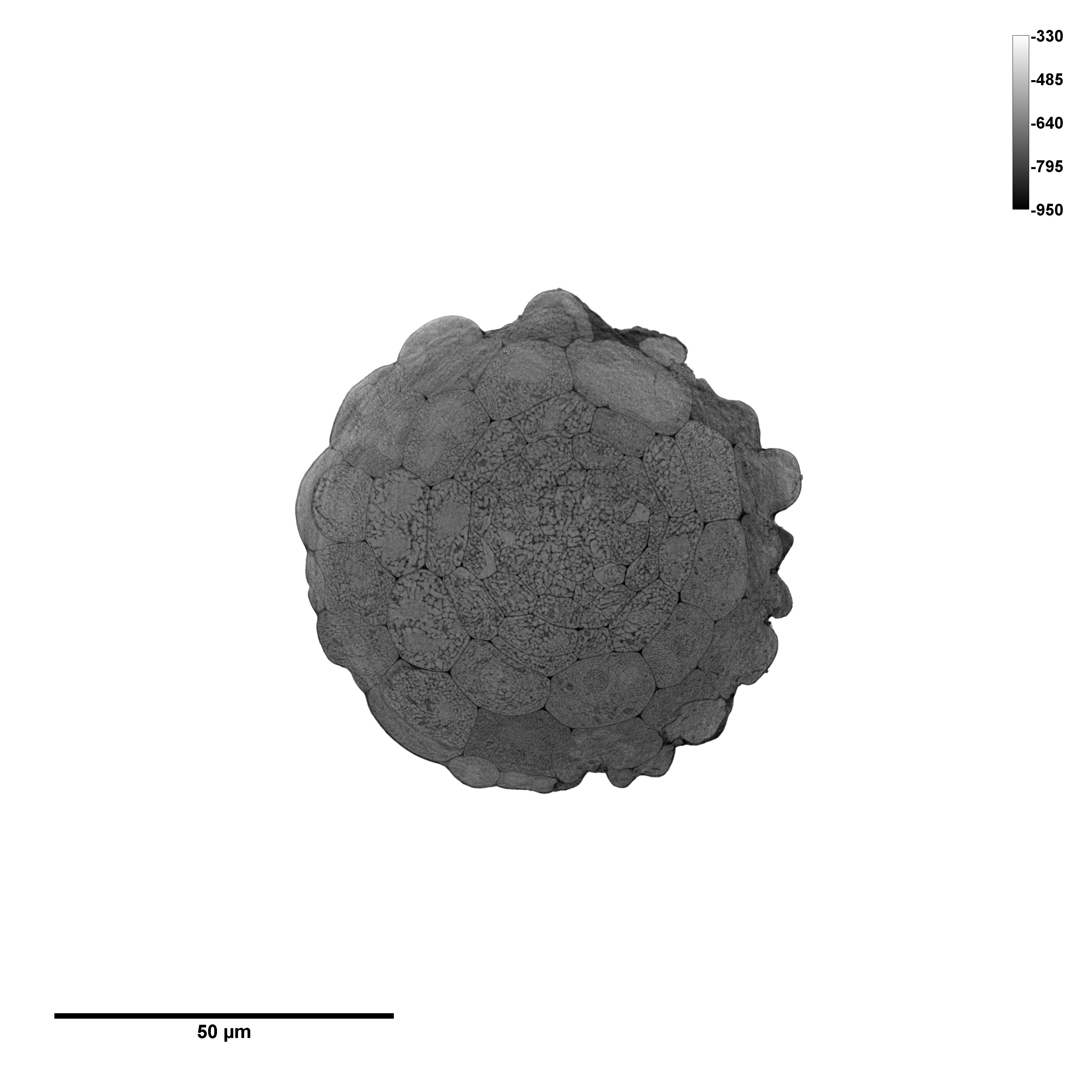

### Se + bact root 3 image 3

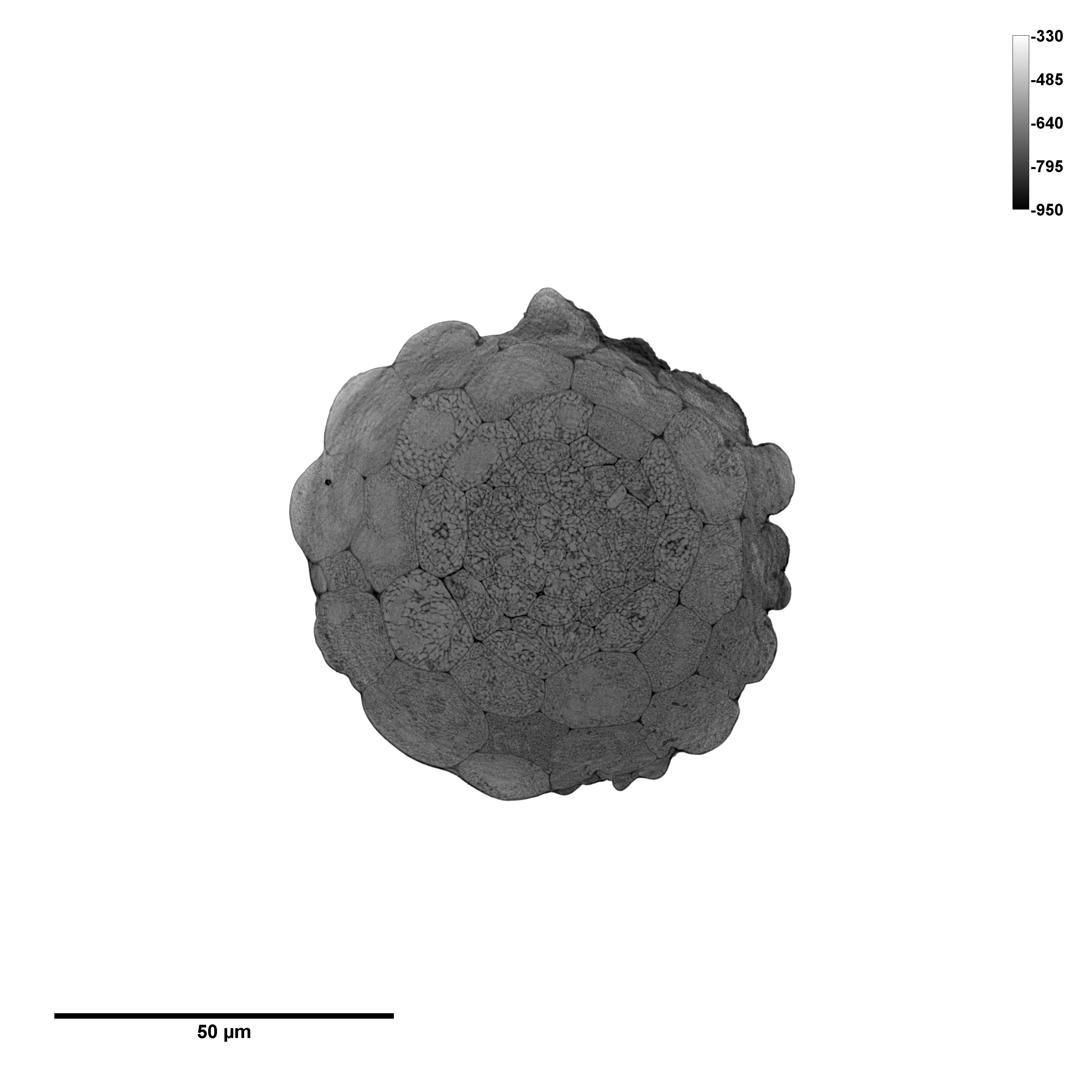

### Se root 1 image 1

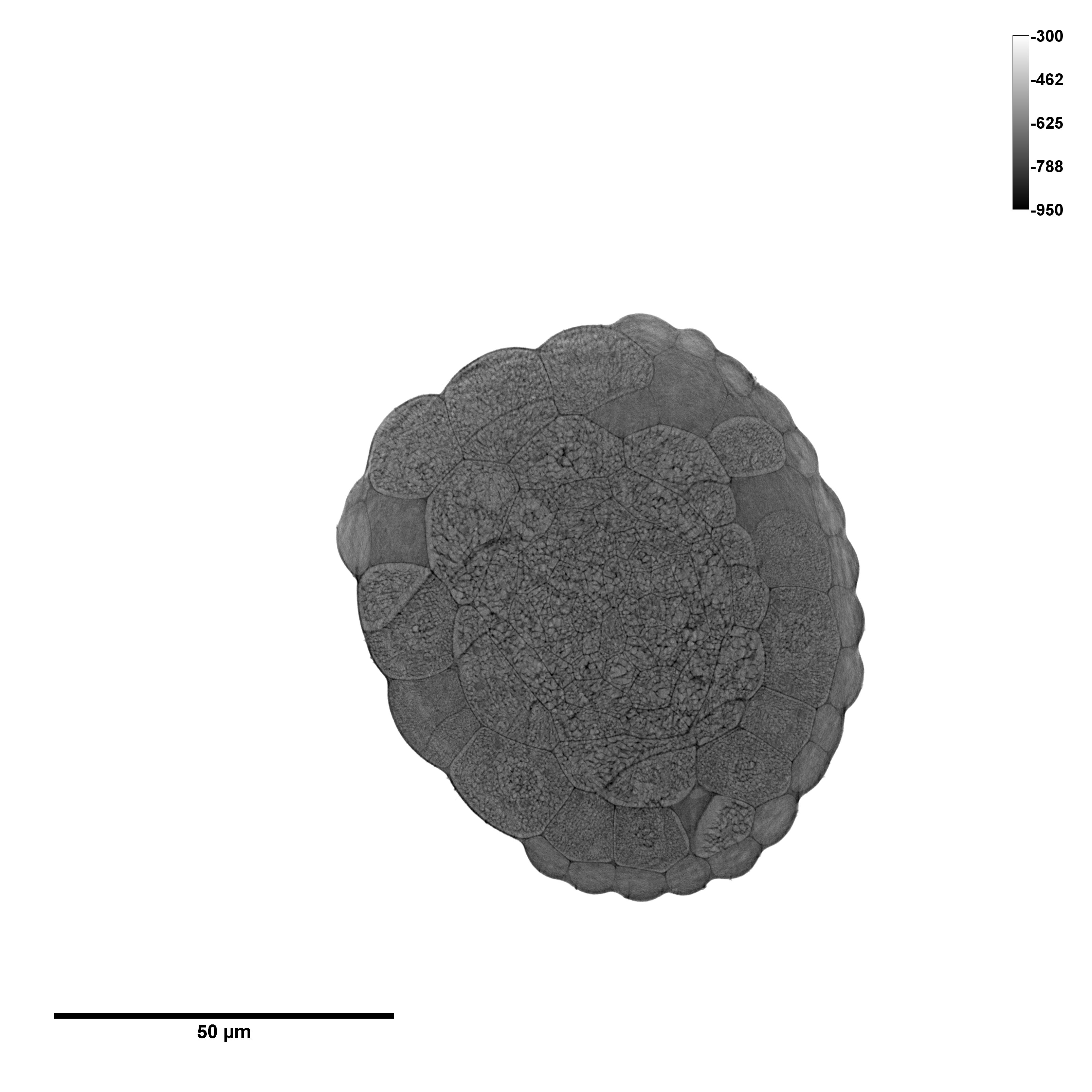

### Se root 2 image 1

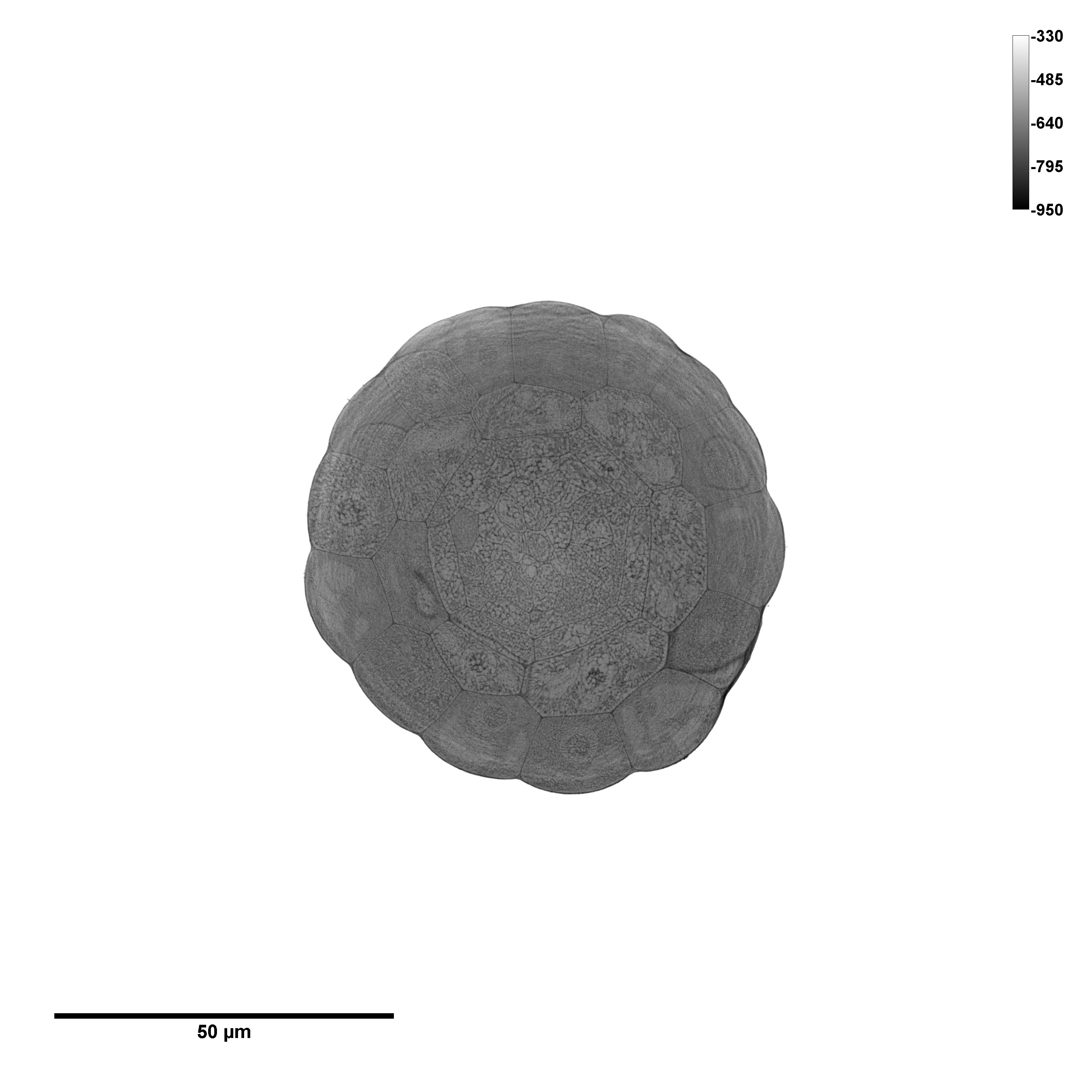

### Se root 2 image 2

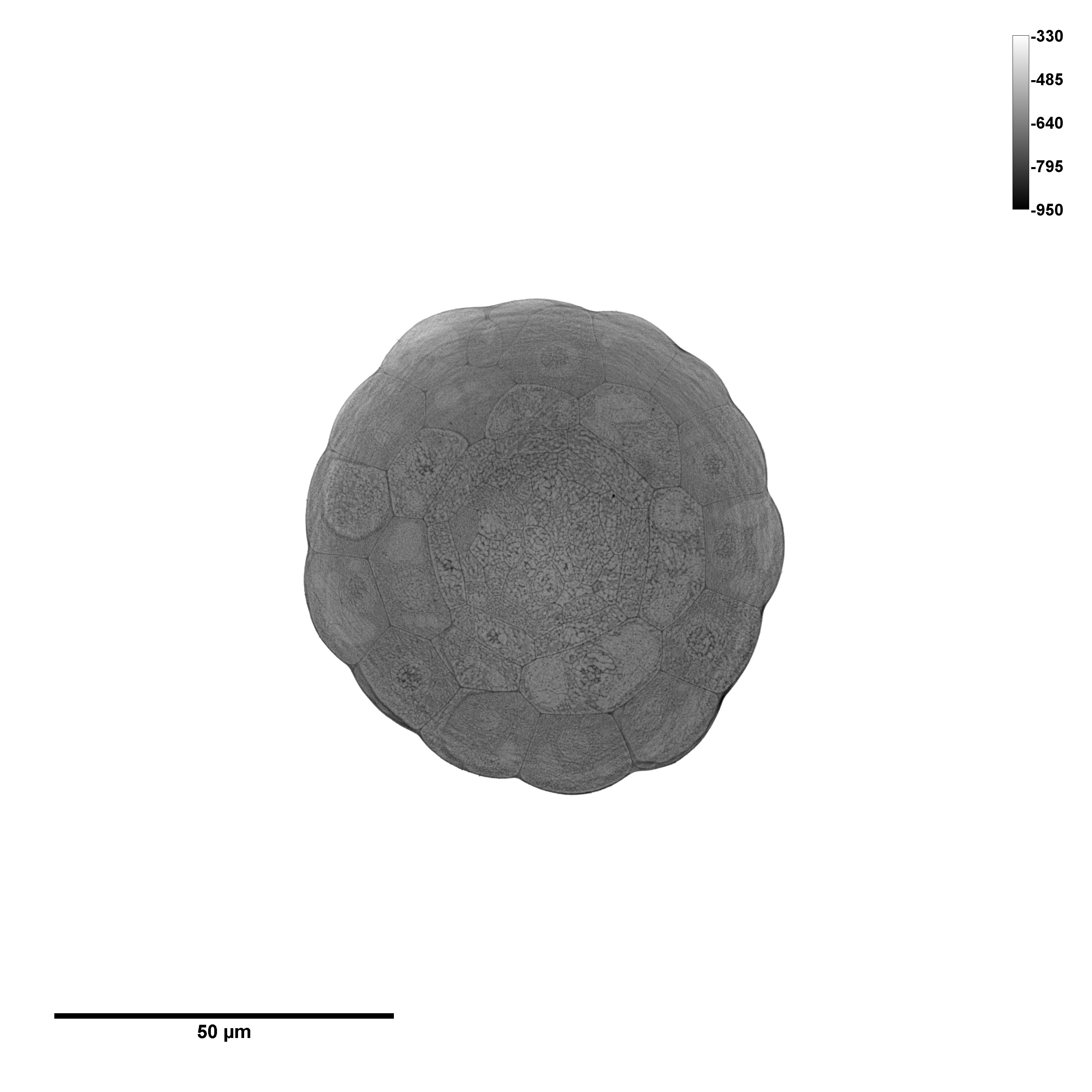

### Se root 3 image 1

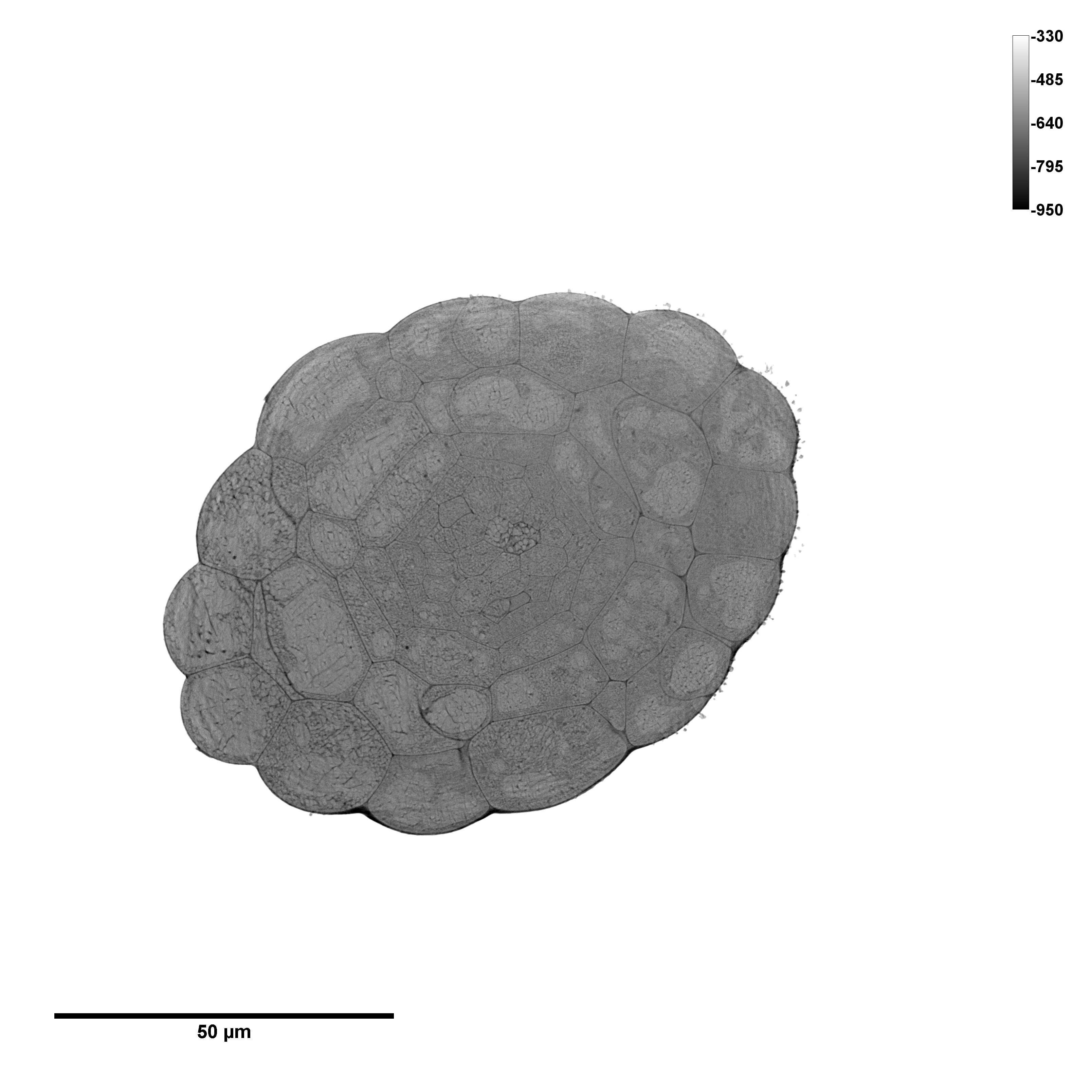
